# Principles for rational Cas13d guide design

**DOI:** 10.1101/2019.12.27.889089

**Authors:** Hans-Hermann Wessels, Alejandro Méndez-Mancilla, Xinyi Guo, Mateusz Legut, Zharko Daniloski, Neville E. Sanjana

## Abstract

Type VI CRISPR enzymes have recently been identified as programmable RNA-guided, RNA-targeting Cas proteins with nuclease activity that allow for specific and robust target gene knock-down without altering the genome. However, we currently lack information about optimal Cas13 guide RNA designs for high target RNA knock-down efficacy. To close this gap, we conducted four massively-parallel Cas13 screens targeting the mRNA of a destabilized green fluorescent protein (GFP) transgene and CD46, CD55 and CD71 cell surface proteins in human cells. In total, we measured the activity of 24,460 guide RNA including 6,469 perfect match guide RNAs and a diverse set of guide RNA variants and permutations with mismatches relative to the target sequences.

We find that guide RNAs show high diversity in knock-down efficiency driven by crRNA-specific features as well as target site context. Moreover, while single mismatches generally reduce knock-down to a modest degree, we identify a critical region spanning spacer nucleotides 15 – 21 that is largely intolerant to target site mismatches. We developed a computational model to identify guide RNAs with high knock-down efficacy. We confirmed the model’s generalizability across a large number of endogenous target mRNAs and show that Cas13 can be used in forward genetic pooled CRISPR-screens to identify essential genes. Using this model, we provide a resource of optimized Cas13 guide RNAs to target all protein-coding transcripts in the human genome, enabling transcriptome-wide forward genetic screens.

---

Type VI CRISPR enzymes have recently been identified as programmable RNA-guided, RNA-targeting Cas proteins with nuclease activity that allow for target gene knock-down without altering the genome. In addition to target RNA knock-down ^1–9^, Cas13 proteins have been used to enable viral RNA detection systems ^7,9–11^, site-directed RNA editing ^12^, demethylation of m^6^A-modified transcripts ^13^, RNA live-imaging ^14,15^, and modulation of splice site choice as well as cleavage and polyadenylation site usage ^5,16,17^.

Cas13 proteins are guided to their target RNAs by a single CRISPR RNA (crRNA) composed of a direct repeat (DR) stem loop and a spacer sequence (guide RNA) that mediates target recognition by RNA-RNA hybridization. Although Cas13 enzymes exert some non-specific collateral nuclease activity upon activation ^4–6,10,18^, they have greatly reduced off-target activity in cultured cells compared to RNA interference ^2,5,12^. Previous studies have shown that Cas13 guide RNAs have minimal Protospacer Flanking Sequence (PFS) constraints in mammalian cells ^1,4,12,19^ and that RNA target sites should be preferentially accessible for Cas13 binding ^1,2,4^. Beyond these basic parameters, we currently lack information about optimal Cas13 crRNA designs for high target RNA knock-down efficacy.

To date, three different Cas13 effector proteins (*Pgu*Cas13b, *Psp*Cas13b, *Rfx*Cas13d) have been reported to show high RNA knock-down efficacy with minimal off-target activity ^5,12^. We compared the ability of these three Cas13 enzymes to knock-down GFP mRNA when directed to either the cytosol or the nucleus. *Rfx*Cas13d (CasRx) consistently showed the strongest target knock-down, especially when fused to a nuclear localization sequence (NLS) (**Supplementary Fig. 1a - c**). Using Cas13d-NLS, we varied the guide length while maintaining a constant guide RNA 5’ end or 3’ end relative to a 30 nt reference guide. In both experiments, we found the most pronounced target knock-down using guide RNAs with a length of 23 – 30 nt (**Supplementary Fig. 1d**). Structural analysis of another Cas13d variant (*Es*Cas13d, PDB: 6E9E/6E9F) suggested that guide RNAs longer than 20 nt extend outside the effector protein binding cleft and that 22 nt guide RNAs provide optimal knock-down ^20^. However, additional guide RNA-target hybridization up to 30 nt in total does not impair target knock-down.

To systematically assess the *Rfx*Cas13d target knock-down efficacy of thousands of guide RNAs, we established a monoclonal HEK293 cell line expressing destabilized GFP and doxycycline-inducible Cas13d protein. We lentivirally delivered a library of 7,500 crRNAs that target the GFP coding sequence, containing perfect match and mismatch guide RNAs (**Fig. 1a**). We performed fluorescence activated cell sorting (FACS) to gate cells in four bins based on their GFP intensity (**Supplementary Fig. 2a**). Guide counts showed high concordance between bins across three independent transductions with clear separation of bin 1, which contained cells with the lowest GFP expression (**Supplementary Fig. 2b-d**).

**Figure 1.**
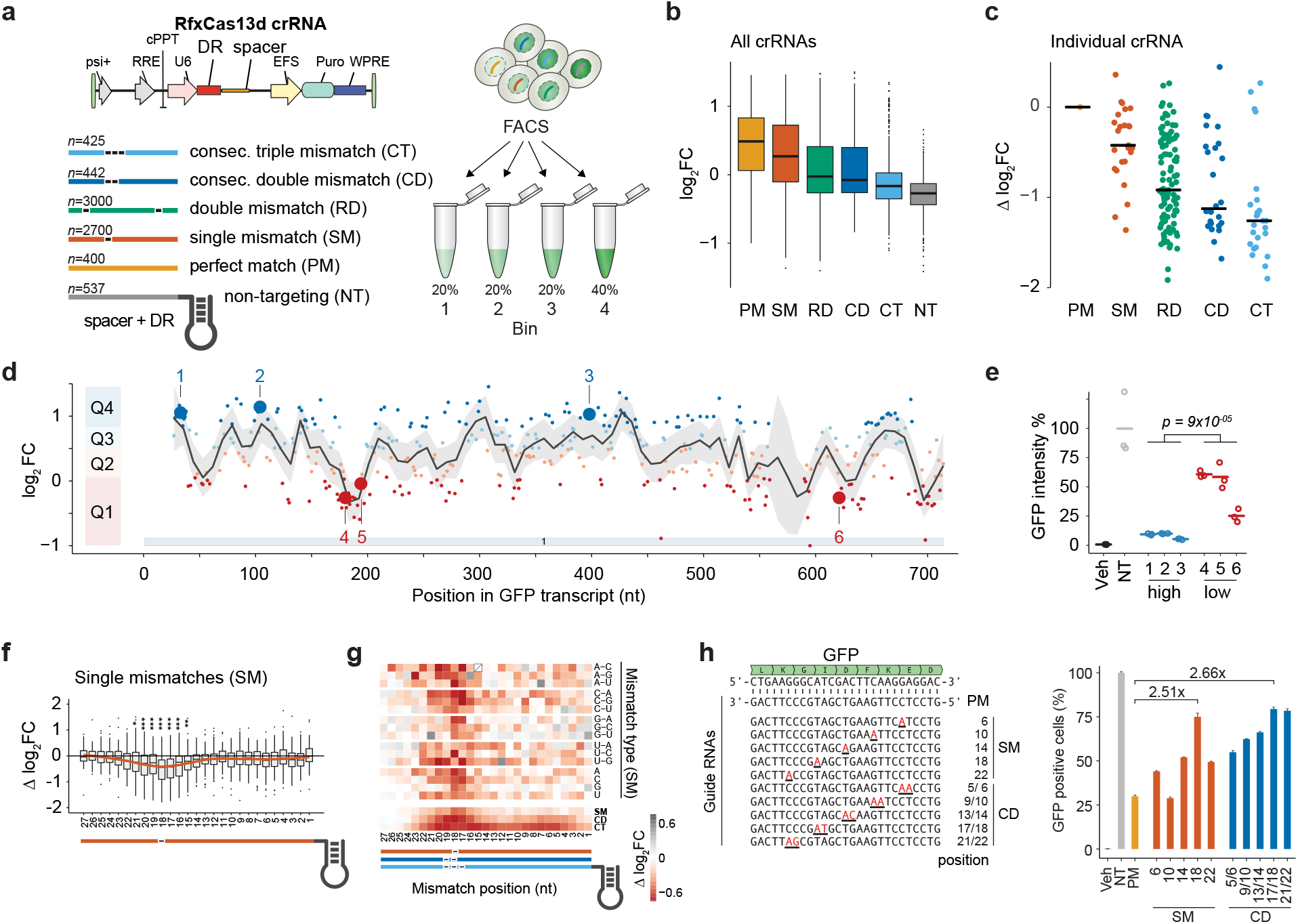
CRISPR Type VI-D *Rfx*Cas13d GFP knock-down pooled tiling screen. (**a**) The GFP-targeting library contained 400 guide RNAs that were perfectly matched, 100 guide RNAs with a single mismatch at each of the 27 guide positions (*n* = 2,700 guides), 30 guide RNAs with 100 random double mismatches (*n* = 3,000 guides), and 17 guides with consecutive double (*n* = 442 guides) and triple (*n* = 425 guides) mismatches. CrRNAs are lentivirally transduced into double-transgenic TetO-*Rfx*Cas13d and GFPd2PEST HEK293 cells. After selection, cells are sorted by GFP intensities into 4 bins. (**b, c**) Log_2_ fold-change (log_2_FC) enrichment scores of guide RNAs comparing guide RNA counts of the lowest fluorescence (Bin 1) to the input (unsorted) cell population. Scores are demarcated by the type of designed crRNAs as given by the list in *a*. (*b*) All crRNAs. (*c*) A single perfect match (PM) crRNA and corresponding derivative crRNAs with mismatches. Guide log_2_FC enrichments are calculated relative to the perfect match reference guide (Δlog_2_FC). Black lines denote medians. (**d**) Distribution of perfect match guide RNAs along the GFP mRNA and their log_2_FC enrichment. Guide RNAs are separated into targeting efficiency quartiles Q1-Q4 with Q4 containing guides with the best knock-down efficiency. (**e**) GFP knock-down validation for 6 guides (3 with high efficacy and 3 with low efficacy) highlighted in *d.* (*n* = 3 transfection replicates; Veh = vehicle transfection, NT = non-targeting crRNAs). Significance from a one-tailed *t*-test. (**f**) Relative targeting efficacy (Δlog_2_FC) of guides with single nucleotide mismatches (SM) at the indicated position relative to their cognate perfect match guides. Significance: * *p* < 0.05, ** *p* < 0.01, *** *p* < 0.001 from a two-tailed *t*-test. (**g**) (*top*) Change in targeting efficacy by guide RNA nucleotide identity or mismatch type. (*bottom*) Change in targeting efficacy for SM, CD, or consecutive triple mismatch (CT) by position. (**h**) Validation of *Rfx*Cas13d seed region. (*left*) Individual perfect match and mismatch guides relative to GFP target mRNA. (*right*) Percent GFP negative cells after co-transfection of specific GFP-targeting crRNAs normalized to the non-targeting control. Veh = vehicle transfection, NT = non-targeting crRNA, PM= perfect match guide RNA.

We calculated the log_2_ fold-change (log_2_FC) crRNA enrichment between all bins and the unsorted input guide RNA distribution (**Supplementary Data 6**). Perfect match guide RNAs were enriched in bin 1, while increasing numbers of mismatches led to a gradual decrease in guide enrichment (**Fig. 1b**, **Supplementary Fig. 3a, b)**. This was true for the whole crRNA population as well as for individual guides and their corresponding guides with 1 – 3 mismatches (**Fig. 1b-c**, **Supplementary Fig. 3c**). As a control, the library also contained 537 non-targeting crRNAs and they were effectively depleted from bin 1 (**Fig. 1b**, **Supplementary Fig. 3a, b**). As expected, guide abundances in bin 1 were negatively correlated to those in bins 2 to 4, which contained cells with higher GFP intensities (**Supplementary Fig. 3d, e**). Taken together, this suggests that the enrichments of guide RNAs in bin 1 accurately reflect target mRNA knock-down.

We noticed considerable heterogeneity of guide enrichment within each class (**Fig. 1b-c**). By examining perfect match guide RNAs along the target mRNA, we observed a position-dependent effect, suggesting an influence of the target sequence context on the guide RNA efficacy (**Fig. 1d**). We selected 6 guides along the GFP target transcript with either high or low enrichment and validated their relative target knock-down efficacies by transfection of individual guides followed by FACS readout (**Fig. 1e**).

To examine if Cas13 can tolerate mismatches between the guide RNA and the target RNA, we calculated the relative log_2_ fold change (Δ log_2_FC) for each mismatch guide by subtracting the log_2_FC from the reference (perfect match) guide (**Fig. 1f**). We found a critical (“seed”) region for Cas13d knock-down efficacy between guide RNA nucleotides 15 to 21 with its center at nucleotide 18 relative to the guide RNA 5’ end. Although seed regions have been shown for Cas13a orthologs ^1,21,22^, one group reported no clear seed region for Cas13d ^20^ while another group showed position-dependent mismatch sensitivity for Cas13d in a cell-free assay ^23^. Within the seed region, single mismatches led to diminished guide enrichment, while mismatches outside the seed region were better tolerated (**Fig. 1f**). The critical region was present irrespective of the mismatch identity (**Fig. 1g**). Similarly, consecutive double and triple mismatches indicated the presence of the critical region (**Fig. 1g**, **Supplementary Fig. 4a**). For randomly distributed double mismatches, the largest change in enrichment was observed in cases where both mismatches are in the seed region (**Supplementary Fig. 4b**). Increasing the number of mismatches to three largely abrogated target knock-down (**Supplementary Fig. 4a**). For this reason, the critical region may have been masked in previous studies on *Rfx*Cas13d which used four consecutive mismatches ^20^.

Given the heterogeneity in enrichment for guide RNAs that have mismatches in the seed region, we sought to assess the effect of surrounding nucleotide context. We used partial correlation to control for the knock-down efficacy of the cognate perfect match guide (“reference”), as poorly performing crRNAs might not allow for large changes in enrichment (**Supplementary Fig. 5a**). Controlling for the reference crRNA efficacy, mismatches in a ‘U’-context in the target site negatively impact Cas13d activity, whereas mismatches in a ‘GC’-context were better tolerated (**Supplementary Fig. 5b**). We confirmed the presence of the seed region in transfection experiments using guides with single or double nucleotide mismatches to the GFP mRNA (**Fig. 1h**). A single mismatch at guide position 18 led to a marked decrease in knock-down efficacy relative to a perfect match guide RNAs. While a perfect match guide decreased the percentage of GFP-positive cells to ~29%, a single mismatch at guide position 18 resulted in 75% GFP-positive cells and a double mismatch at positions 17 and 18 resulted in ~79% GFP positive cells (**Fig. 1h**).

Importantly, the center of the *Rfx*Cas13d seed region coincides with conserved contacts between a helical domain in Cas13d protein and the backbone of the guide RNA-target hybrid interface. This interaction resides opposite of the guide RNA position 17-18 with the target RNA ^20^. The helical domain is located between both HEPN-domains needed for target cleavage, and mutation of the interacting amino acids in *Es*Cas13d completely abolished target cleavage ^20^. Mismatches at the seed center thus may impair HEPN-domain activity.

Next we sought to assess the features that may affect knock-down efficacy for perfect match guide RNAs (see **Supplementary Note 1** for details). One of the features impacting the observed guide RNA enrichments in the GFP tiling screen was crRNA folding: Predicting secondary structures and corresponding minimum free energy (MFE) of perfect match crRNAs showed a positive correlation between the MFE and guide efficacy (**Supplementary Fig. 6a**). In particular, ‘G’-dependent structures, such as predicted G-quadruplexes, showed diminished target knock-down. Given that the crRNA folding is critical for effective target knock-down, we sought to further stabilize and improve the DR by repairing a predicted bulge in the DR, by varying the length of the stem loop or by disrupting bases in the proximal DR stem (**Supplementary Fig. 6b**). Analysis of the crystal structure of *Es*Cas13d and *Ur*Cas13d together with its crRNA suggested that the terminal loop in the DR may not be embedded within the protein and thus may allow extension (and further stabilization) of the stem loop ^20,23^ similar to those previously found to enhance Cas9 activity or utility ^24,25^. We observed that any change in stem length abrogated target knock-down completely (**Supplementary Fig. 6c**). Also, repairing the bulged nucleotide within the stem loop decreased target knock-down. However, disrupting the first base pair within the proximal stem further increased Cas13d targeting efficacy, leading to a novel *Rfx*Cas13d DR with improved knock-down capability. We tested the modified DR on 6 additional guides targeting GFP and found that the modified DR improved target knock-down especially for guides with low knock-down efficiency (**Supplementary Fig. 6d**).

We defined 15 crRNA and target-RNA features based on their correlation with observed guide enrichment in our exploratory data analysis (**Supplementary Table 1, Supplementary Note 1, Supplementary Data 7**). With these features, we sought to derive a generalizable ‘on-target’ model to predict Cas13d target knock-down. We compared the ability of machine learning approaches to predict guide efficiency (observed log_2_FC) in the held-out data (see Methods) and found that a Random Forest (RF) model had the best prediction accuracy (**Supplementary Fig. 7a**), weighting the crRNA folding energy, the local target ‘C’-context, and the upstream target ‘U’-context as the most important features (**Supplementary Fig. 7b**). Other learning approaches frequently chose similar features, suggesting that these features are the main drivers of Cas13d GFP knock-down (**Supplementary Fig. 7c**). To identify key predictor of guide efficiency, we iteratively reduced the number of features, monitoring the model performance and derived a minimal model that explained about 37% of the variance (*r*^*2*^) with a Spearman correlation (*r*_*s*_) of ~0.58 to the held-out data (**Fig. 2a**, **Supplementary Fig. 7d**). In comparison, an support vector machine (SVM) regression model with a similar structure to a Cas9 guide prediction algorithm ^26^ performed worse when applied to this data (*r*^*2*^ = 0.21, *r*_*s*_ = 0.44) (**Fig. 2a**). We used 10-fold cross-validation to confirm that the model can readily separate poor performing guide RNAs from effective crRNAs. Accordingly, 46% of the guides present in the highest efficacy-quartile are predicted to reside in the best performing quartile. Conversely, 64% of guides present in the lowest efficacy-quartile are predicted to reside in the poorest performing quartile (**Supplementary Fig. 7e**). Similarly, the predicted standardized guide score of the *N* top- or bottom-ranked crRNAs confirmed that the model can effectively separate crRNAs that perform well from those that perform poorly (**Supplementary Fig. 7f**).

**Figure 2.**
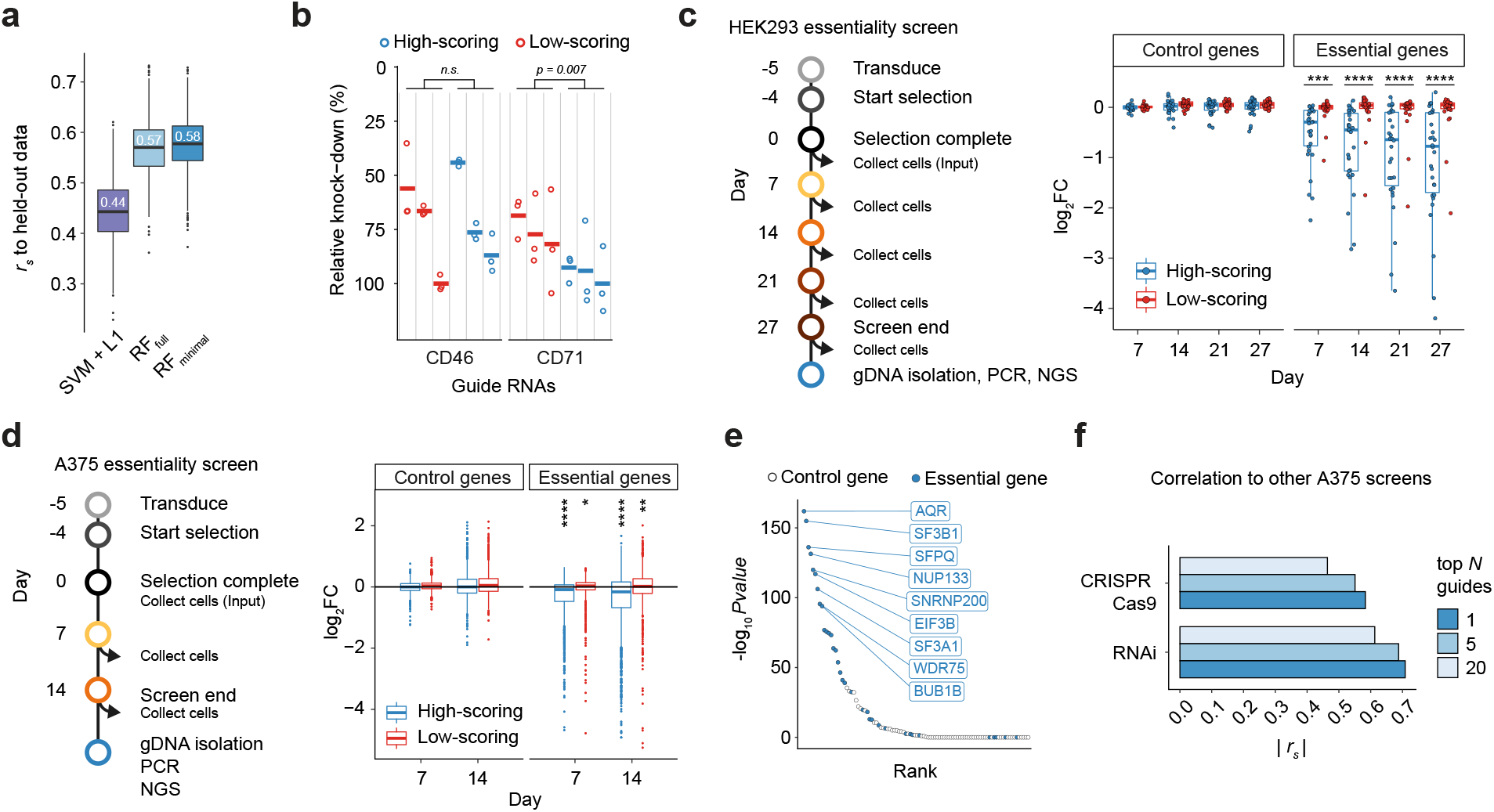
*Rfx*Cas13d on-target guide RNA prediction model. (**a**) Correlation of predictions from a Random Forest (RF) regression model (either with all features or a minimal set of the most predictive features) and a support vector machine with L1 regression to held-out screen data. (**b**) Validation of on-target model testing 3 high-scoring and 3 low-scoring guide RNAs via targeting of cell-surface proteins and antibody labeling to measure target knock-down by FACS. Relative knock-down indicates the percent reduction (relative to non-targeting guide RNAs) in the mean fluorescence intensity. (*n* = 3 transfection replicates). (**c**) Validation of on-target model assaying 3 high-scoring and 3 low-scoring guide RNAs per gene in a gene essentiality screen in HEK293 cells with growth dropout phenotype testing 10 essential genes and 10 control genes. Each point represents one guide as a mean of three replicate experiments. The y-axis depicts the log_2_ fold-change (FC) of the guide RNA at the indicated time point relative to the Day 0 sample. One-sided KS-test comparing high-scoring and low-scoring guides, *** *p* = 2×10^−5^, **** *p* = 2×10^−6^. (**d**) A375 essentiality screen with growth dropout phenotype assaying 20 high-scoring and 20 low-scoring guide RNAs per gene. One-sided KS-test comparing high-scoring or low-scoring guides to the distribution of non-targeting guides. * *p* = 0.043, ** *p* = 0.0095, **** *p* < 1×10^−44^. (**e**) Gene ranking for essentiality based on the <u>r</u>obust <u>r</u>ank <u>a</u>ggregation (RRA) *p*-value across replicates for all 20 high scoring guides. Blue dots denote essential genes from a prior RNAi screen ^27^. (**f**) Spearman rank correlation of Cas13d gene depletion (as in *e*) with prior CRISPR-Cas9 and RNAi screens in A375 cells. Analysis includes genes represented in all libraries (*n* = 35 essential genes and *n* = 15 control genes). (RNAi screen: A375 DEMETER2 v5 score ^27^, Cas9 screen: A375 STARS score ^28^).

To show that our model is generalizable, we predicted guides to target the endogenous transcripts of *CD46* and *CD71*, which encode cell surface proteins, and measured the guide knock-down efficacy by FACS. For each gene, we chose 3 guide RNAs predicted to have high knock-down efficacy (Q3 or Q4) and 3 guide RNAs predicted to have low knock-down efficacy (Q1 or Q2). On an individual guide level, we found that the majority of guides with higher predicted guide scores suppressed CD46 and CD71 protein expression more robustly than guides with lower guide scores (**Fig. 2b**). Comparing the observed knock-down across all three high-scoring to all three low-scoring guide RNAs, we found a significant improvement for CD71, while for CD46 we observed considerable variance. To increase throughput and test guide RNA efficacy predictions for more genes, we first generated a small crRNA library targeting 10 essential and 10 control genes with both 3 high-scoring and 3 low-scoring guide RNAs and monitored their depletion in a gene essentiality screen over time. Essential genes were chosen from genes that were strongly depleted in previous RNAi screens ^27^ (**Supplementary Fig. 8a**). Most high-scoring guides targeting essential genes were progressively depleted over time, while low-scoring guides showed largely no depletion (**Fig. 2c**, **Supplementary Fig. 8b**).

In addition, we performed a second targeted essentiality screen in A375 cells targeting 35 essential and 65 control genes with both 20 high-scoring and 20 low-scoring guide RNAs. Similar to the HEK293 screen above, we found that high-scoring guides that target essential genes were progressively depleted over time (**Fig. 2d**, **Supplementary Fig. 8c**). Although high-scoring guide RNAs were generally more depleted than low-scoring guide RNAs on a per gene level, we noticed that not all predicted essential genes showed depletion upon Cas13d targeting (**Supplementary Fig. 8c-d**), suggesting that RNAi-screen derived essentiality scores may not be one-to-one transferable to Cas13d derived essentiality.

We calculated a significance score of gene depletion based on the guide rank consistency for the 20 high-scoring guides and found strong enrichment of defined essential genes at the top of the list (**Fig. 2e**). The guide RNA depletion scores correlated better with the DEMETER2 RNAi ^27^ scores used to define the set of essential genes to be tested (up to *r*_*s*_ = 0.71 using the best guide) than with the Cas9 STARS scores ^28^ (up to *r*_*s*_ = 0.61) (**Fig. 2f**). Taken together, this suggest that the crRNA and target RNA features derived from the GFP tiling screen can generalize to predict Cas13d guide efficacy for novel targets, and that these guide predictions can be used in pooled CRISPR-Cas13 screens.

Our predictive on-target model based on the GFP tiling screen was largely able to separate guide RNAs with low knock-down efficiency from those with high efficiency. However, given that we observed remaining heterogeneity among the predicted high-scoring guides, we sought to improve our on-target model by enlarging our training dataset. Therefore, we performed three additional Cas13d tiling screens targeting the main transcript isoforms of the cell surface proteins CD46, CD55 and CD71 in HEK293 cells coupled with FACS readout selecting for cells with decreased surface protein expression (**Fig. 3a-c**; **Supplementary Fig. 9a-b**). Besides perfect match guide RNAs, we added several guide RNA classes (**Supplementary Fig. 9a**). For each screen, perfect match guide RNAs showed the strongest guide enrichment relative to the unsorted input samples, while reverse complement negative control guides and non-targeting guides were depleted (**Supplementary Fig. 9c**). In the new screens we reduced the overall guide length to 23 bases and included a set of guide length variants ranging in length from 15 to 36 nucleotides. Starting from 23 nucleotide length, guides RNAs exerted full knock-down efficiency, while longer guide 3’ends did not have any deleterious effects (**Supplementary Fig. 9d**).

**Figure 3.**
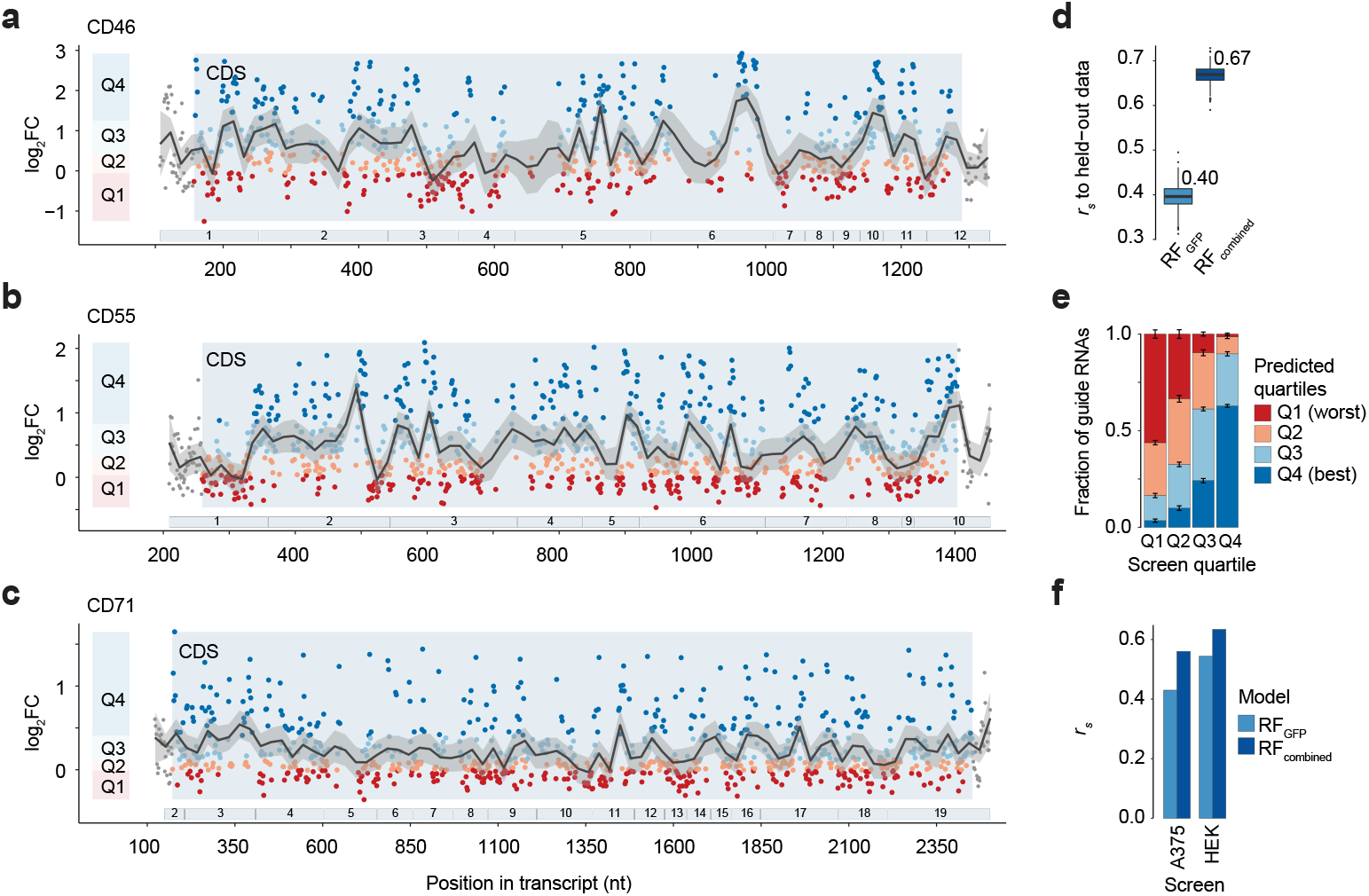
Improvement of *Rfx*Cas13d on-target guide RNA prediction model with tiling screens over endogenous transcripts. (**a-c**) Distribution of perfect match guide RNAs along the coding region (CDS) of CD46, CD55 and CD71 mRNA and their log_2_ fold-change (FC) enrichments. Positive FC values indicate better transcript knock-down. Guide RNAs are separated into targeting efficiency quartiles Q1-Q4 per gene with Q4 containing guides with the best knock-down efficiency. Numbered bars below indicate exons. (**d**) Correlation of predictions from the RF_minimal_ (=RF_GFP_) model and the updated RF_combined_ regression model to held-out screen data using bootstrapping across all four tiling screens. (**e**) Comparison of predicted and measured log_2_FC quartiles across the 10-fold model cross-validation. Quartile definition as in a-c. (**f**) Spearman rank correlation between observed guide RNA depletion (= target knock-down) and the predicted guide score for the indicated Cas13d essentiality screens and indicated on-target models (see **Fig 2c-d** and **Supplementary Fig. 8**).

Perfect match guides targeting coding regions (CDS) were more strongly enriched compared to guides targeting untranslated regions (UTRs) or introns (**Supplementary Fig. 9e**). UTR-targeting guides may show lower enrichments as each target gene may be represented by multiple transcript isoforms with alternative UTR usage. Hence, guides targeting coding regions have a higher likelihood to find the cognate target site while, for example, 3’UTR-targeting guide RNAs find their target site only in a fraction of the expressed transcripts isoforms. Accordingly, the low enrichment for intron-targeting guide RNAs may be explained by the short-lived nature of introns. For these guides, the intronic target site is present only for a short period of time, which likely enables the transcript to evade Cas13 targeting. For this reason, guide RNA knock-down efficiency may not be directly comparable between CDS-targeting guides and UTR- or intron-targeting guides.

We also observed a slight decrease in guide efficiency of intron-targeting guides immediately downstream of the 5’-splice-site and within the −50 to 0 nucleotide upstream of the 3’-splice-site summarizing across all 39 introns present (**Supplementary Fig. 9f**). These sites are typically bound by the spliceosome ^29^, suggesting that guide RNAs targeting these regions may compete with the splice machinery and other splice factors for the target sequences. As transcript maturation in the nucleus seemingly influences the guide RNA targeting efficiency, we wondered if the exon-junction-complex (EJC) would affect knock-down of the matured transcript in the same way. The EJC typically binds ~20-24 nucleotides 5’ upstream to the exon-exon-junction upon splicing ^30,31^. Indeed, we observed a depletion of high-scoring guide RNAs within a window of −20 to 0 nucleotides 5’ upstream to the exon junction (**Supplementary Fig. 9g**).

To improve our on-target model, we focused on perfect match guide RNAs that target CDS-regions and increased the number of high-confidence model input observation from ~400 to nearly 3000. Similar to the initial GFP-screen, guide RNAs efficiencies were distributed along the coding region in a non-random manner (**Fig. 3a-c**). We repeated the assessment of features that may affect knock-down efficacy (see **Supplementary Note 2** for details). Notably, the increased number of observations uncovered positional nucleotide preferences (**Supplementary Fig. 10a-b**). Guide enrichments correlated positively with G- and C-base probabilities in the seed region around guide position 18. And surrounding this region U- and A-base probabilities correlate positively with the target knock-down. We derived an updated on-target model using 2,918 CDS-targeting guide RNAs across all four tiling screens and selected 35 out of 644 evaluated features in a similar fashion as before (see methods) (**Supplementary Table 2, Supplementary Note 2, Supplementary Data 7**).

The combined Random Forest model (RF_combined_) displayed improved prediction accuracy compared to the initial RF_minimal_ model (from here on referred to as RF_GFP_) explaining ~47% of the variance (*r*^*2*^) with a Spearman correlation (*r*_*s*_) of ~0.67 to the held-out data (**Fig. 3d**, **Supplementary Fig. 10c**). Using 10-fold cross-validation the model effectively separated low-scoring guides from high-scoring guides, assigning 63% of the guide RNAs correctly to the highest efficacy-quartile (**Fig. 3e**). Similarly, the predicted guide scores of the top- or bottom-ranked guide RNAs (ranked by the observed knock-down efficiency) separate guides that performed well from those that performed poorly more than expected by chance (**Supplementary Fig. 10d**). Further, we performed leave-one-out cross-validation training on three data sets while predicting guide scores for the held-out fourth screen. The RF_combined_ model generalized well for endogenous genes (mean ± sd: *r*_*s*_ = 0.63 ± 0.01) but was less predictive for the GFP transgene (*r*_*s*_ = 0.33) (**Supplementary Fig. 10e**).

Finally, we compared the ability of both models, the RF_GFP_ and RF_combined_ model, with respect to their ability to correctly predict the knockdown efficiencies for the two essentiality screens. Both screens were designed based on guide predictions made by the RF_GFP_ model. In both cases, the RF_combined_ was in better agreement with the observed knock-down efficiencies across all genes (**Fig. 3f**). Likewise, we found that the RF_combined_ showed improved agreement with the observed guide RNA depletion also on a gene level for the 10 most depleted genes in the A375 fitness screen (RF_GFP_: *r*_*s*_ = 0.46 ± 0.16, RF_combined_: 0.58 ± 0.14). Taken together, we show that our updated RF_combined_ on-target model is able to predict Cas13d guide RNA target knock-down efficiencies, separating poorly performing guides from guides with high efficacy and generalized across numerous targets.

We applied our model and predicted guide RNAs for all protein-coding transcripts in the human genome (GENCODE v19). We made these predictions available through a user-friendly, web-based application (https://cas13design.nygenome.org). In addition, we report the 10 highest-scoring crRNAs for the 5’ UTR, CDS and 3’ UTR of each transcript (**Supplementary Fig. 11a, Supplementary Data 8**). We partitioned the predicted guide RNAs according to the efficacy quartiles in our four screens. Only 15.2% of all possible guides fall into the highest scoring (best knock-down) quartile (Q4) (**Supplementary Fig. 11b)**. A large fraction of guide RNAs are predicted to have lower efficacy (36.8% of all guides are in Q1 or Q2), which emphasizes the value of optimal guide selection for high knock-down efficacy. However, almost all transcripts have top-scoring guide predictions (**Supplementary Fig. 11c**).

Taken together, we performed a set of pooled screens for CRISPR Type VI Cas13d and defined targeting rules for optimal guide design. We show that crRNA choice and target RNA-context constrain target knock-down efficacy and, using this data, we develop and validate an ‘on-target’ model to predict guides with high efficacy. Although we specifically sought to define rules for active Cas13d, we believe that our model may be transferable to inactive (catalytically dead) Cas13d effector proteins. Beyond our on-target guide design, we identified a critical seed region in the crRNA that is sensitive to target mismatch. We provide evidence that this seed region can be used in living cells to discriminate between target RNAs with high similarity, such as allele-specific single nucleotide polymorphisms.

## Methods

### Cloning of Cas13 nuclease, guide RNAs and destabilized EGFP plasmids

Using Gibson cloning, we modified the EF1a-short (EFS) promoter-driven lentiCRISPRv2 (Addgene 52961) or lentiCas9-Blast (Addgene 52962) plasmids with several different transgenes^32^. For the destabilized EGFP construct, we introduced a PEST sequence and nuclear localization tag on EGFP to create EFS-EGFPd2PEST-2A-Hygro (pLentiEGFPdestabilized) from lentiCas9-Blast. To test the upstream U-content, we introduced a multiple cloning site (MCS) into pLentiEGFPdestabilized right after the stop codon, and used the MCS to introduce oligonucleotide sequences with variable U-content^32^.

For the CRISPR Type-VI orthologs, we cloned effector proteins (*Pgu*Cas13b: Addgene 103861, *Psp*Cas13b: Addgene 103862, *Rfx*Cas13d: Addgene 109049) and their direct repeat (DR) sequences (*Pgu*Cas13b: Addgene 103853, *Psp*Cas13b: Addgene 103854, *Rfx*Cas13d: Addgene 109053) into lentiCRISPRv2. In this manner, we created pLentiRNACRISPR constructs: hU6-[Cas13 DR]-EFS-[Cas13 ortholog]-[NLS/NES]-2A-Puro- WPRE, where [Cas13 ortholog] was one of *Pgu*Cas13b, *Psp*Cas13b, or *Rfx*Cas13d and [NLS/NES] was either a nuclear localization signal or nuclear export signal. To generate doxycycline-inducible Cas13d cell lines, we cloned NLS-*Rfx*Cas13d-NLS (Addgene 109049) into TetO-[Cas13]-WPRE-EFS-rtTA3-2A-Blast. For the screens, we changed the DR in the lentiGuide-Puro vector (Addgene 52963) to contain the *Rfx*Cas13d DR using Gibson cloning to create lentiRfxGuide-Puro (pLentiRNAGuide) ^32^. All plasmids will be made available on Addgene.

Guide cloning was done as described previously^32^. All constructs were confirmed by Sanger sequencing. All primers used for molecular cloning and guide sequences are shown in **Supplementary Data 1**.

### Cell culture and monoclonal cell line generation

HEK293FT cells were acquired from Thermo Fisher Scientific (R70007) and A375 cells were acquired from ATCC (CRL-1619). HEK293FT and A375 cells were maintained at 37°C with 5% CO_2_ in D10 media: DMEM with high glucose and stabilized L-glutamine (Caisson DML23) supplemented with 10% fetal bovine serum (Serum Plus II Sigma-Aldrich 14009C) and no antibiotics.

To generate doxycycline-inducible *Rfx*Cas13d-NLS HEK293FT and A375 cells, we transduced cells with a *Rfx*Cas13d-expressing lentivirus at low MOI (<0.1) and selected with 5μg/mL Blasticidin S (ThermoFisher A1113903). Single cell colonies were picked after by sparse plating. Clones were screened for Cas13d expression by western blot using mouse anti-FLAG M2 antibody (Sigma F1804).

For the GFP tiling screen *Rfx*Cas13d-expressing cells were transduced with pLentiEGFPdestabilized lentivirus at low MOI (<0.1) and selected with 100μg/ml Hygromycin B (ThermoFisher 10687010) for 2 days. Single-cell colonies were grown by sparse plating. Resistant and GFP-positive clonal cells were expanded and screened for homogenous GFP expression by FACS.

### Transfection and flow cytometry

For all transfection experiments, we seeded 2×10^5^ HEK293FT cells per well of a 24-well plate prior to transfection (12 - 18 hours) and used 500 or 750 ng plasmid together with a 5-to-1 ratio of Lipofectamine 2000 (ThermoFisher 11668019) or 1 mg/mL polyethylenimine (Polysciences 23966) to DNA (e.g. 2.5μl Lipofectamine2000 or PEI mixed with 0.5μg plasmid DNA). Flow cytometry or fluorescence-assisted cell sorting (FACS) was performed at 48 hrs post-transfection. All transfection experiments were performed in biological triplicate.

For the CRISPR Type-VI ortholog comparison (**Supplementary Fig. 1a-c**), we cloned the effector proteins (*Pgu*Cas13b: Addgene 103861, *Psp*Cas13b: Addgene 103862, *Rfx*Cas13d: Addgene 109049) and their direct repeat sequences (*Pgu*Cas13b: Addgene 103853, *Psp*Cas13b: Addgene 103854, *Rfx*Cas13d: Addgene 109053) as described above. We co-transfected the pLentiRNACRISPR constructs together with a GFP expression plasmid in a 2:1 molar ratio. The guide RNA length comparison (**Supplementary Fig. 1d**) was done using previously published *Rfx*Cas13d constructs (Addgene 109049 and 109053), except that we removed the GFP cassette from the *Rfx*Cas13d plasmid. The modified *Rfx*Cas13d construct and guide plasmids were co-transfected together with a GFP expression plasmid in a 2:2:1 molar ratio. For the DR modification experiment (**Supplementary Fig. 6c**) we transfected *Rfx*Cas13d expressing cells, starting doxycycline-induction (1μg/ml) at the time of cell plating. The guide plasmid and GFP expression plasmid were co-transfected at a 1:1 molar ratio.

For the model validation flow cytometry (**Fig. 2b**) we transfected *Rfx*Cas13d-expressing cells with a guide RNA expressing plasmid. 48 hours post transfection, the cells were stained for the respective cell surface protein for 30 min at 4°C and measured by FACS. (BioLegend: CD46 #352405 clone TRA-2-10, CD71 (TFRC) #334105 clone CYIG4).

For the screen result validation (**Fig. 1e**) and seed validation experiments (**Fig. 1h**) we co-transfected *Rfx*Cas13d-expressing cells with a guide RNA expressing plasmid and GFP plasmid at a 1:1 molar ratio. At 48 hours post-transfection, the cells were analyzed by flow cytometry.

To assess the upstream U-context (**Supplementary Note 1**), we transfected upstream-U context modified pLentiEGFPdestabilized-MCS plasmid together with either a crRNA plasmid into *Rfx*Cas13d-expressing in a 2:1 molar ratio. Each GFP-upstreamU-context plasmid was co-transfected with both a targeting or a non-targeting guide RNA used for calculating the knock-down, as a change in 3’UTR uridine content could attract RNA-binding proteins that may affect RNA stability independent of Cas13. We selected the zero-uridine oligonucleotide from a set of 10000 *in silico* randomized 52mers with {A_24_,C_14_,G_14_} with minimal predicted RNA-secondary structure as determined by RNAfold ^33^ with default setting.

For flow cytometry analysis, cells were gated by forward and side scatter and signal intensity to remove potential multiplets. If present, cells were additionally gated with a live-dead staining (LIVE/DEAD Fixable Violet Dead Cell Stain Kit, Thermo Fisher L34963). For each sample we analyzed at least 5000 cells. If cell numbers varied, we randomly sampled all samples to the same number of cells before calculating the mean fluorescence intensity (MFI). For GFP co-transfection experiments, we only considered the percentage of transfected cells with the highest GFP expression determined by comparing the non-targeting control to wild-type control cells. For the upstream U-context co-transfection experiments, we considered the whole cell populations.

For knock-down experiments of endogenous genes (**Fig. 2b**), we determined the percentage of transfected cells with lower target gene signal than the non-targeting control in the condition with the highest observed knock-down. For all conditions, we analyzed the same bottom percentage of cells. For the selected cells, we compared the MFI of targeting guides relative to non-targeting guides to determine the percent knock-down. To directly compare relative rank of individual guides as done in **Fig. 2b**, we normalized the effect size by setting the most effective guide to 100%. For the seed validation (**Fig. 1f**), we determined the percentage of transfected (GFP-positive) cells with GFP signal higher than Lipofectamine vehicle treated control cells. The percentage of transfected cells was normalized to percentage of GFP-positive cells in the non-targeting guide control.

### Screen library design and pooled oligo cloning

To design the *Rfx*Cas13d guide RNA library for GFP, we selected the 714 bp coding sequence (without start codon) to be targeted. *In silico*, we generated all perfectly matching 27mer guide RNAs with minimal constraints (T-homopolymer < 4, V-homopolymer < 5, 0.1 < GC-content < 0.9) and selected 400 by random sampling. From these, we sampled 100 guide RNAs and introduced one random nucleotide conversion at each position (*n* = 2700, SM set). From these 100, we randomly sampled 17 guide RNAs and introduced 26 or 25 consecutive double (*n* = 442, CD set) and triple (*n* = 425, CT set) mismatches, respectively. We sampled an additional 13 guide RNAs from the SM set (in total, 30 guide RNAs) and introduced 100 random double mismatches at any position for each guide RNA if not present already in the set of 17 consecutive double mismatches (*n* = 3000, RD set). In total, we designed 6,967 GFP targeting guides and added 533 non-targeting guides (NT set) of the same length from randomly generated sequences that did not align to the human genome (hg19) with less than 3 mismatches.

For CD46, CD55 and CD71 library design, we selected the transcript isoform with highest isoform expression in HEK-TE samples (determined by Cancer Cell Line Encyclopedia CCLE; GENCODE v19) and longest 3’UTR isoform (CD46: ENST00000367042.1, CD55: ENST00000367064.3, CD71: ENST00000360110.4). As described above, we generated all perfectly matching 23mers, and selected ~2000 evenly spaced guide RNAs per target. In addition to PM, SM, RD and NT sets as described above, we included for each target a set of guide length variants (*n* = 450, LV set), guide RNAs targeting intronic sequences near splice-donor and splice-acceptor sites across all 39 annotated introns (*n* = 2122, I set) and an additional negative control set of reverse complementary perfect match sequences (*n* = 300, RC set). Further details are in **Supplementary Data 2**.

For both targeted essentiality screens, we used the DEMETER2 v5 ^27^ data set from the Cancer Dependency Map portal (DepMap) to determined essential and control genes. Specifically, we selected essential genes with low log_2_ fold-change (FC) enrichments across all cell lines and in the respective assay cell line (**Supplementary Fig. 8a,c**). For our HEK293FT cells, we considered data for HEK-TE cells. Furthermore, we selected genes with one transcript isoform constituting more than 75% of the gene expression with expression level less than ~150 transcripts per million (TPM). We predicted guide RNA efficiencies using the minimal RF_GFP_ model and removed all guides with matches or partial matches elsewhere in the transcriptome. We allowed up to 3 mismatches when looking for potential off-targets. From the set of remaining perfect match guide RNA predictions, we manually selected three high-scoring and three low-scoring guides for the HEK293FT cell line screen to ensure that each guide fell into non-overlapping regions of the target transcripts. For the A375 cell line targets, we selected the top 20 high-scoring guide RNAs. For the set of 20 low-scoring guides, we chose among the bottom 60 to reduce the overlap of guide RNAs that fall into the same region. In this way, we assayed 20 genes in HEK293FT cells targeting 10 essential and 10 control genes with three low-scoring and three high-scoring guides, as well as three non-targeting guides (*n* = 123). For the A375 screen, we targeted 100 genes (35 essential and 65 control genes) with 40 guides each (20 high- and 20 low-scoring) and included 680 non-targeting sequences (*n* = 4680).

All large-scaled pooled crRNA libraries were synthesized as single-stranded oligonucleotides (Twist Biosciences), PCR amplified using NEBNext High-Fidelity 2X PCR Master Mix (M0541S) (**Supplementary Data 1**), and Gibson cloned into pLentiRfxGuide-Puro. The guides for the HEK293FT essentiality screen were ordered from IDT, array cloned, confirmed by Sanger sequencing, and subsequently pooled using equal amounts. Complete library representation with minimal bias (90^th^ percentile/10^th^ percentile crRNA read ratio: 1.68 – 2.17) were verified by Illumina sequencing (MiSeq).

### Pooled lentiviral production and screening

Lentivirus was produced via transfection of library plasmid with appropriate packaging plasmids (psPAX2: Addgene 12260; pMD2.G: Addgene 12259) using polyethylenimine (PEI) reagent in HEK293FT. At 3 days post-transfection, viral supernatant was collected and passed through a 0.45 um filter and stored at −80C until use.

Doxycycline-inducible *Rfx*Cas13d-NLS human HEK293FT, double-transgenic HEK293FT-GFP or A375 cells were transduced with the respective library pooled lentiviruses in separate infection replicates ensuring at least 1000x guide representation in the selected cell pool per infection replicate using a standard spinfection protocol. We generated either 2 or 3 independent replicate experiments. After 24 hours, *Rfx*Cas13d expression was induced by addition of 1μg/ml doxycycline (Sigma D9891) and cells were selected with 1 ug/mL puromycin (ThermoFisher A1113803), resulting in ~30% cell survival. Puromycin-selection was complete ~48 post puromycin-addition. Assuming independent infection events (Poisson), we determined that ~83% of surviving cells received a single sgRNA construct. Cells were passaged every two days maintaining at least the initial cell representation and supplemented with fresh doxycycline.

The tiling screens were terminated after 5 to 10 days. For all targets we noted maximal knock-down after 2-4 days (data not shown). For cell surface proteins, cells were stained in batches of 1×10^7^ cells for 30 min at 4°C (BioLegend: CD46 clone TRA-2-10 #352405 - 3μl per 1×10^6^ cells; CD55 clone JS11 #311311 - 1.5μg per 1×10^6^ cells; CD71 clone CYIG4 #334105 - 4μl per 1×10^6^ cells). We collected unsorted samples for input guide RNA representation of approximately 1000x coverage for each sample and sorted at least another 1000x representation into the assigned bins based on their signal intensities (GFP: lowest 20%, 20%, 20% and remaining highest 40%, **Supplementary Fig. 2a**; CD proteins lowest 20% and highest 20%, **Supplementary Fig. 9b**; **Supplementary Data 2**). Cells were PBS-washed and frozen at −80°C until sequencing library preparation. In each case, the bin containing the lowest 20% represented the strongest target knock-down.

The essentiality screens were started (Day 0) upon complete puromycin selection, which was at 5 days after transduction. Cells were passaged every two to three days maintaining at least the initial cell representation and supplemented with fresh doxycycline. At Day 0 (=Input) and every 7 days, we collected a >1000x representation from each sample. The HEK293FT cell screen was conducted in triplicate and cultured for 4 weeks. The A375 cell screen was conducted in duplicate and cultured for 2 weeks.

### Screen readout and read analysis

For each sample, genomic DNA was isolated from sorted cell pellets using the GeneJET Genomic DNA Purification Kit (ThermoFisher K0722) using 2×10^6^ cells or less per column. The crRNA readout was performed using two rounds of PCR ^34^. For the first PCR step, a region containing the crRNA cassette in the lentiviral genomic integrant was amplified from extracted genomic DNA using the PCR1 primers in **Supplementary Data 1**.

For each sample, we performed PCR1 reactions as follows: 20 μl volume with 2 ug of gDNA in each reaction limited by the amount of extracted gDNA (total gDNA ranged from 8μg to 50 ug per sample with an estimated representation of 10^6^ diploid cells per ~6.6 ug gDNA. PCR1: 4μl 5x Q5 buffer, 0.02U/μl Q5 enzyme (M0491L), 0.5uM forward and reverse primers and 100ng gDNA/μl. PCR conditions: 98°C/30s, 24x[98°C/10s, 55°C/30s, 72°C/45s], 72°C/5min).

We pooled the unpurified PCR1 products and used the mixture for a single second PCR reaction per sample. This second PCR adds on Illumina sequencing adaptors, barcodes and stagger sequences to prevent monotemplate sequencing issues. Complete sequences of the 5 forward and 3 reverse Illumina PCR2 readout primers used are shown in **Supplementary Data 1**. (PCR2: 50μl 2x Q5 master mix (NEB #M0492S), 10μl PCR1-product, 0.5uM forward and reverse PCR2-primers in 100μl. PCR conditions: 98°C/30s, 17x[98°C/10s, 63°C/30s, 72°C/45s], 72°C/5min).

Amplicons from the second PCR were pooled by screen experiment (e.g. all GFP-screen samples) in equimolar ratios (by gel-based band densitometry quantification) and then purified using a QiaQuick PCR Purification kit (Qiagen 28104). Purified products were loaded onto a 2% E-gel and gel extracted using a QiaQuick Gel Extraction kit (Qiagen 28704). The molarity of the gel-extracted PCR product was quantified using KAPA library quant (KK4824) and sequenced on an Illumina NextSeq 500 - II MidOutput 1×150 v2.5.

Reads were demultiplexed based on Illumina i7 barcodes present in PCR2 reverse primers using bcl2fastq and by their custom in-read i5 barcode using a custom python script. Reads were trimmed to the expected guide RNA length by searching for known anchor sequences relative to the guide sequence using a custom python script. For the tiling screens, pre-processed reads were either aligned to the designed crRNA reference using bowtie ^35^ (v.1.1.2) with parameters `-v 0 -m 1` or collapsed (FASTX-Toolkit) to count perfect duplicates followed by string-match intersection with the reference to retain only perfectly matching and unique alignments. Pre-processed guide RNA sequences from the essentiality screens were aligned allowing for up to 1 mismatch (`-v 1 -m 1`).Alignment statistics are available in **Supplementary Data 3**. The raw guide RNA counts (**Supplementary Data 4**) were normalized separated by screen dataset using a median of ratios method like in DESeq2 ^36^ and underwent batch-correction using combat implemented in the SVA R package ^37^. Non-reproducible technical outliers were removed by applying pair-wise linear regression for each sample after normalization and batch-correction, collecting the residuals and taking the median value for each guide RNA across all sample-centric comparisons. We removed all crRNA counts within the top *X*% residuals across all samples (GFP: 2%, CD proteins: 0.5%, Essentiality screen: no outlier removal). For the GFP screen, we only remove outliers on a per-sample basis as needed (but not the entire guide RNA). For CD46, CD55 and CD71 screens, since the number of outliers was small, we decided to remove the entire guide RNA from the analysis. The table below indicates all filtering applied:

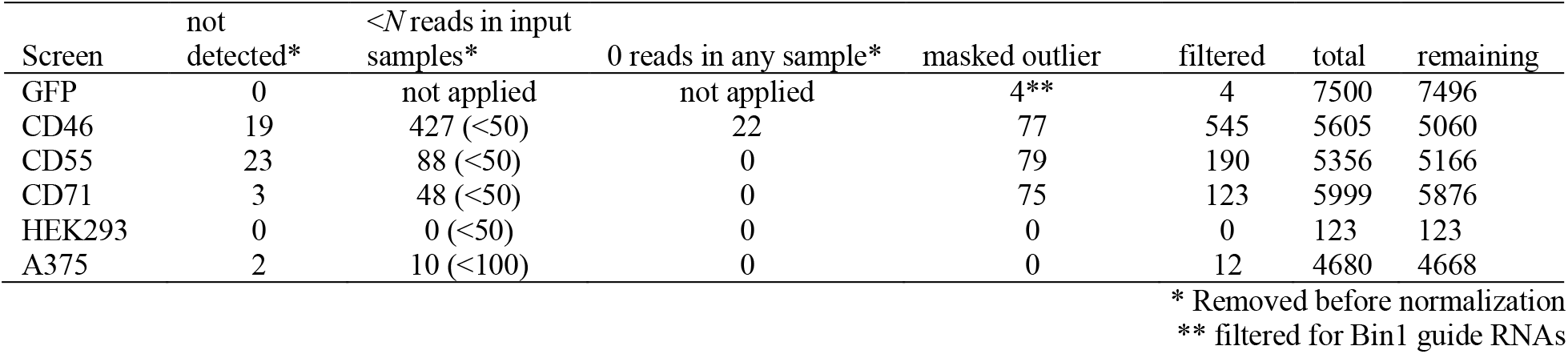

| Screen | not detected* | <N reads in input samples* | 0 reads in any sample* | masked outlier | filtered | total | remaining |
| --- | --- | --- | --- | --- | --- | --- | --- |
| GFP | 0 | not applied | not applied | 4** | 4 | 7500 | 7496 |
| CD46 | 19 | 427 (<50) | 22 | 77 | 545 | 5605 | 5060 |
| CD55 | 23 | 88 (<50) | 0 | 79 | 190 | 5356 | 5166 |
| CD71 | 3 | 48 (<50) | 0 | 75 | 123 | 5999 | 5876 |
| HEK293 | 0 | 0 (<50) | 0 | 0 | 0 | 123 | 123 |
| A375 | 2 | 10 (<100) | 0 | 0 | 12 | 4680 | 4668 |

\* Removed before normalization
\*\* filtered for Bin1 guide RNAs

Processed crRNA counts are available in **Supplementary Data 5**. Guide RNA enrichments were calculated building the count ratios between a bin or timepoint and the corresponding input sample and log_2_-transformation (log_2_FC). Consistency between replicates was estimated using robust rank aggregation (RRA) ^38^. Delta log_2_FC for mismatching guides was calculated by subtracting the log_2_FC of the perfectly matching reference guide. For the tiling screens, all plots and analyses were performed using the mean guide RNA enrichments of bin 1 (= bottom 20%) across replicates, unless indicated otherwise. Similarly, we used the mean guide RNA enrichments relative to Day 0 across replicates for the essentiality screen. Guide RNA enrichment scores (log_2_FC) are available in **Supplementary Data 6.** In all combined analyses across all four tiling screens, we scaled the observed log_2_FC separately to improve comparability. For the generation of a the combined on-target model, we normalized the 2918 selected CDS-targeting guides RNA across the four tiling screens to the same scale prior to training and testing the model. To do so, for each dataset *D*, we computed the upper and lower quartiles of the guide log_2_FC (*UQ*_*D*_ and *LQ*_*D*_, respectively) as well as the corresponding quartiles for the log_2_FC among all datasets pooled together (*UQ*_*P*_ and *LQ*_*P*_). We then updated each fold change *x* as follows: *x̂* =[(*x* − *LQ*_*D*_) / (*UQ*_*D*_ − *LQ*_*D*_) * (*UQ*_*P*_ − *LQ*_*P*_) + *LQ*_*P*_]. By centering on quartiles, this procedure normalized the fold-change distributions in a way that was less susceptible to the influence of outliers of a single screen.

### Predicting RNA secondary structures and RNA-RNA hybridization energies

crRNA secondary structure and minimum free energies (MFEs) was derived using RNAfold [`--gquad`] on the full-length crRNA (DR + guide) sequence ^33^. For building the combined on-target model and for testing the RF_GFP_ model on the combined data set, we assumed 23mer guide RNAs for all guides in the GFP tiling screen to prevent length dependent differences in the crRNA MFE. Target RNA unpaired probability (accessibility) was calculated using RNAplfold [`-L 40 -W 80 -u 50`] as described before ^39^. We performed a grid-search calculating the RNA accessibility for each target nucleotide in a window of minus 20 bases downstream of the target site to plus 20 bases upstream of the target site assessing the unpaired probability of each nucleotide over 1 to 50 bases for all perfectly matching guides. Then, we calculated the Pearson correlation coefficient between the log_10_-transformed unpaired probabilities and the observed guide RNA log_2_FC for each point and window relative to the guide RNA.RNA-RNA-hybridization between the guide RNA and its target site was calculated using RNAhybrid [`-s -c`] ^40^. For the hybridization calculation, we did not include the direct repeat of the crRNA. We calculated the RNA-hybridization minimum free energy for each guide RNA nucleotide position *p* over the distance *d* to the position *p* + *d* with its cognate target sequence. All measures were either directly correlated with the observed crRNA log_2_FC or using partial correlation to account for the crRNA folding MFE. In each case, we computed the Pearson correlation.

### Assessing guide RNA nucleotide composition

Guide RNA composition was derived by calculating the nucleotide probability within the respective guide RNA sequence length. To assess the presence of sequence constraints similar to a previously described anti-tag^19^ or 5’ and 3’ Protospacer Flanking Sequences (PFS), we ranked all perfectly matching guide RNAs by their log_2_FC enrichment within each screen separately. We selected the top and bottom 20% enriched/depleted guide RNAs and calculated the positional nucleotide probability for the four nucleotides upstream and downstream relative to the guide RNA match. To assess nucleotide preferences at any guide RNA match position in addition to upstream and downstream nucleotides, we selected the top 20% of the log_2_FC-ranked perfectly matching guides as described above and calculated nucleotide preferences as described before ^26^. In brief, we calculated the probability of each nucleotide at each position for the top guide RNAs and all guide RNAs. The effect size is the difference of nucleotide probability by subtracting the values from all guides from the top guides (delta log_2_FC). *p*-values were calculated from the binomial distribution with a baseline probability estimated from the full-length GFP mRNA target sequence for all perfectly matching crRNAs. *p*-values were adjusted using a Bonferroni multiple hypothesis testing correction.

### Assessing target RNA context

To assess the target RNA context, we calculated the nucleotide probability at each position (*p*) over a window (*w*) of 1 to 50 nucleotides centered around the position of interest (e.g. *p* = −18 with *w* = 11 summarizes the nucleotide content in a window from −23 to −13 with +1 being the first base of the crRNA). We evaluated *p* for all positions within 75 nucleotides upstream and downstream of the guide RNA. The nucleotide context of each point was then correlated with the observed log_2_FC crRNA enrichments for all perfect match crRNAs, either directly or using partial correlation accounting for crRNA folding MFE. In each case we used Pearson correlation.

The RNA context around single nucleotide mismatches was assessed accordingly with a slight modification. Here, the nucleotide context was assessed relative to mismatch position summarizing the nucleotide probability in a window of 1 to 15 nucleotides to either side (e.g. *p* = 18 with *w* = 5 summarizes the nucleotide content in a window of 11 nucleotides from 23 to 13). For more details on *p* and *w*, please see the diagram in **Supplementary Figure 5b**. We used all 2,700 single nucleotide mismatch guides in the GFP tiling screen (100 guide RNAs x 27 mismatched positions per guide). The nucleotide context of each position and each window size was then correlated with the observed delta log_2_FC relative to the perfectly matching reference guide RNA, either directly or using partial correlation accounting for crRNA folding MFE. In each case, we used Pearson correlation.

### On-target model selection

An explanation for all selected features for the RF_GFP_ and RF_combined_ model can be found in **Supplementary Table 1** and **Supplementary Table 2**, respectively. The RF_combined_ model feature input values can be found in **Supplementary Data 7**. All continuous feature scores were scaled to the [0, 1] interval limited to the 5^th^ and 95^th^ percentile, with a mean set to the 5^th^ percentile. Scaled values exceeding the [0, 1] interval were set to 0 or 1, respectively.

Scaling parameters used to normalize data to the [0, 1] interval for the Random Forest Models

| **Model** | **Feature** | **5<sup>th</sup> percentile** | **95<sup>th</sup> percentile** |
| --- | --- | --- | --- |
| **RF<sub>GFP</sub>** | crRNA MFE | -23.4000 | -14.5000 |
|  | Local A probability | 0.0000 | 0.4286 |
|  | Local C | 0.2273 | 0.5000 |
|  | Local G probability | 0.1429 | 0.4286 |
|  | Local U probability | 0.0556 | 0.2778 |
|  | upstream U probability | 0.0667 | 0.2000 |
| **RF<sub>combined</sub>** | crRNA MFE | -20.3000 | -12.8000 |
|  | Log<sub>10</sub> Unpaired probability | -7.5546 | -1.6244 |
|  | hybridization MFE nt 3-15 | -29.4000 | -17.9000 |
|  | hybridization MFE nt 15-23 | -21.8000 | -12.3000 |
|  | Local A<sub>max</sub> probability | 0.0000 | 0.5000 |
|  | Local C<sub>max</sub> probability | 0.0000 | 0.5000 |
|  | Local G<sub>max</sub> probability | 0.0000 | 0.6667 |
|  | Local U<sub>max</sub> probability | 0.0833 | 0.5000 |
|  | Local AU<sub>max</sub> probability | 0.2727 | 0.7272 |
|  | Local GC<sub>max</sub> probability | 0.2222 | 0.7778 |
|  | Local A<sub>min</sub> probability | 0.0000 | 0.5714 |
|  | Local C<sub>min</sub> probability | 0.0000 | 0.5556 |
|  | Local G<sub>min</sub> probability | 0.0000 | 0.4444 |
|  | Local U<sub>min</sub> probability | 0.0000 | 0.5000 |
|  | Local AU<sub>min</sub> probability | 0.2222 | 0.7778 |
|  | Guide A nt probability | 0.0870 | 0.4348 |
|  | Guide C nt probability | 0.0870 | 0.3913 |
|  | Guide G nt probability | 0.0870 | 0.4348 |
|  | Guide AA di-nt probability | 0.0000 | 0.1818 |
|  | Guide AC di-nt probability | 0.0000 | 0.1364 |
|  | Guide AG di-nt probability | 0.0000 | 0.1818 |
|  | Guide AU di-nt probability | 0.0000 | 0.1364 |
|  | Guide CA di-nt probability | 0.0000 | 0.1818 |
|  | Guide CC di-nt probability | 0.0000 | 0.1364 |
|  | Guide CG di-nt probability | 0.0000 | 0.1364 |
|  | Guide CU di-nt probability | 0.0000 | 0.1364 |
|  | Guide GA di-nt probability | 0.0000 | 0.1364 |
|  | Guide GC di-nt probability | 0.0000 | 0.1818 |
|  | Guide GG di-nt probability | 0.0000 | 0.1818 |
|  | Guide GU di-nt probability | 0.0000 | 0.1364 |
|  | Guide UA di-nt probability | 0.0000 | 0.1364 |
|  | Guide UC di-nt probability | 0.0000 | 0.1818 |
|  | Guide UG di-nt probability | 0.0000 | 0.1818 |

To evaluate and compare model performances, we randomly sampled 1,000 bootstrap datasets from the data of perfect match guide RNA log_2_FC response values and selected features. We used 399 data points for the initial RF_GFP_ model and 2918 data points for all CDS-annotating perfect match guides across the four tiling screens. For the RF_combined_ model we normalized the observed log_2_FC values data prior to training and testing as described earlier. Normalized response values showed better generalizability compared to unnormalized or scaled log_2_FC. For each bootstrap sample, 70% of the data was used for training and the remaining 30% of the data was held out for testing, ensuring a 70/30 split for each screen dataset when testing the RF_combined_ model. Linear dependencies between features were identified using the function findLinearCombos from the R package caret and removed. The model performance was evaluated by calculating the Spearman correlation coefficient *r*_*s*_ and Pearson *r*^2^ to the held-out data. We compared a variety of different methods ^39^ (see Table below).

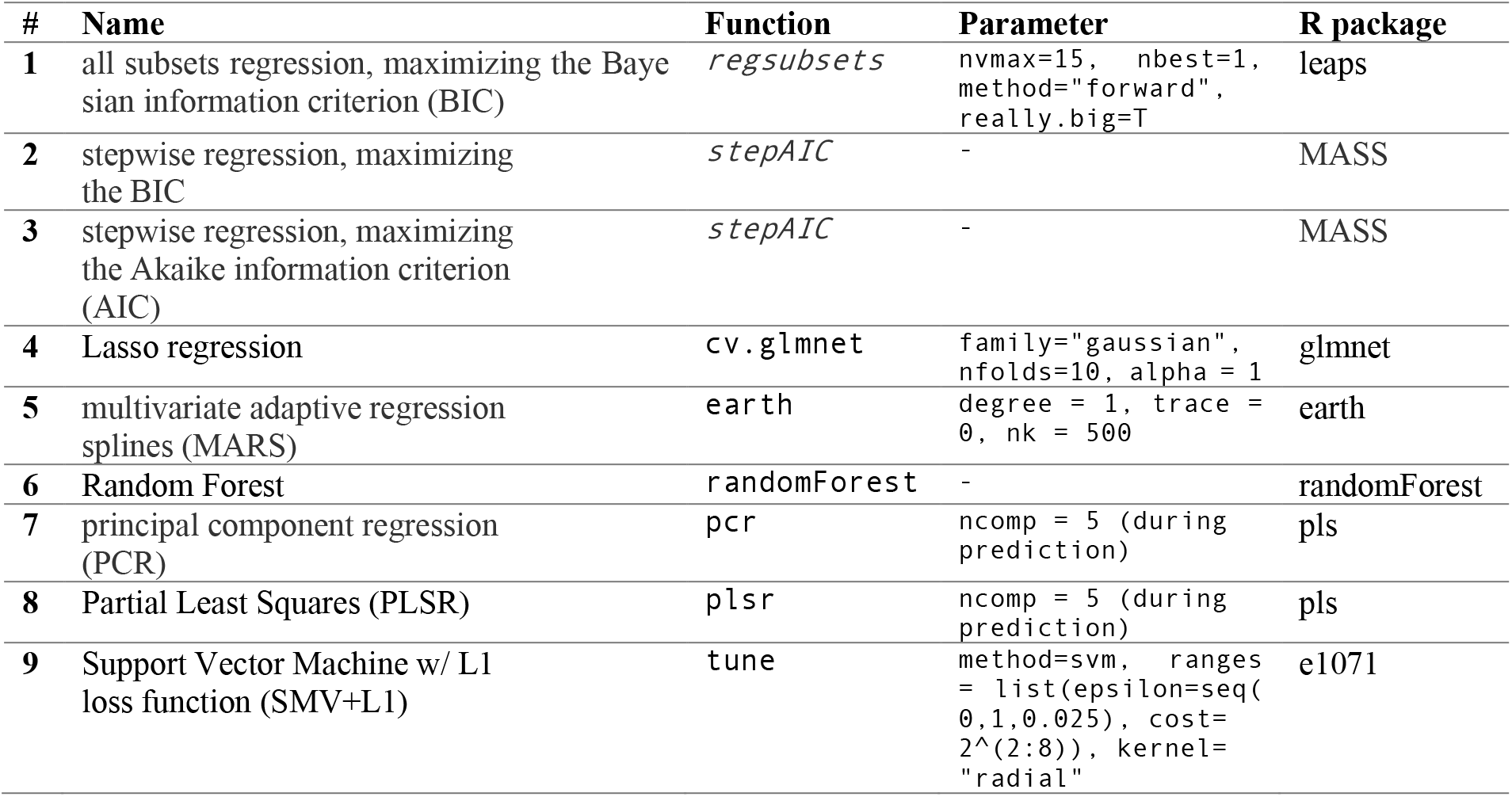

For both models, we tested a variety of feature combinations including crRNA folding energies, RNA-RNA hybridization energies, target site accessibility, overall and positional (di-)nucleotide probabilities, and one-hot encoding for single and di-nucleotide of the guide target-sites and their upstream and downstream flanking four nucleotides. Together, these represented 644 features for the combined on-target model. A full set of features for the combined on-target model can be found in **Supplementary Data 7**. For the initial on target model based on the GFP screen data, we evaluate a set of 15 defined features (**Supplementary Table 1**) along-side with one-hot encoded positional nucleotide information and GC content. These 15 features were defined based on their positive or negative correlation to the observed response value during the data exploration (see also **Supplementary Note 1**). We iteratively reduced the numbers of features from 15 to 6 for the RF_GFP_ model and monitored the model performance as described above. At each iteration, the Random Forest model performed slightly better than any other learning approach. Reducing the features to fewer than the selected 6 features (RF_minimal_ = RF_GFP_) reduced the model performance. For the combined on-target model, we did not iteratively reduce the set of 35 selected features. We compared the RF_GFP_ model to an SVM+L1 model similar to one of the first CRISPR-Cas9 on-target model. Specifically, we used one-hot encoding for all 35 nucleotide positions considered (27 guide RNA positions and 8 additional positions with 4 upstream and 4 downstream nucleotides). Considering all positions, the feature space contained 140 single nucleotide features, 544 di-nucleotide features and the GC-content (685 non-all-zero features). Here, we used tuning (see table below for parameters) to increase model performance for SVM+L1 specifically. Here, but also for the combined model, one-hot encoded features did not lead to high Spearman correlation coefficient *r*_*s*_ to the held-out data.

For further evaluation of the random forest models we used 10-fold cross-validation by randomly partitioning the data into 10 equally-sized partitions ensuring even contribution from each screen to each partition. We trained the model 10 times on 90% of the data and predicted the held-out 10%. For each data point, we assigned the known guide RNA efficacy quartile based on the log_2_FC enrichment and compared it the predicted efficacy quartiles in the held-out data. We also assessed the predicted guide score by calculating the median predicted guide score for the top and bottom ranked crRNAs in the 10% held-out data based on the known log_2_FC-rank for all 10 cross-validation folds (top/bottom *N* = 2, 4, 8, 16, 32, 64, 128 or 256 guide RNAs). To compute the null distribution, we calculated the median predicted guides scores of randomly selected guide RNAs across 1000 samplings for each *N*. For the leave-one-out cross-validation we trained on all data from three tiling screens and performed Spearman rank correlation of the predicted the guide efficiency of the held-out fourth screen to the observed log_2_FC enrichments.

To make the guide score more interpretable, we standardized the guide score to a [0, 1] interval preserving the distribution between 5^th^ and 95^th^ percentile. Normalized values exceeding the [0, 1] interval were set to 0 or 1, respectively. The final RF_GFP_ model was trained on all data points for perfect match guides using the six selected features with 1500 regression trees. The model explains 36.9% of the observed variance with a mean of squared residuals of 0.139. The table below shows the feature contribution for the RF_GFP_ model.

| Feature | %IncMSE | IncNodePurity |
| --- | --- | --- |
| crRNA MFE | 57.989 | 22.617 |
| Local A probability | 30.542 | 7.529 |
| Local C probability | 46.255 | 13.683 |
| Local G probability | 38.557 | 9.256 |
| Local U probability | 29.953 | 6.555 |
| upstream U probability | 31.559 | 8.629 |

Similarly, final RF_combined_ was trained on 2918 data points using 35 selected features. Tuning the number of trees (`ntree`) and number of splitting variables per node (`mtry`) led to insignificant insignificant performance improvements compared to default settings. The model (`mtry` = 12, `ntree` = 2000) explains 47.16% of the observed variance, a mean of squared residuals of 0.168, and the feature contribution as indicated below ranked by importance:

| Feature | %IncMSE | IncNodePurity |
| --- | --- | --- |
| Local G<sub>max</sub> probability | 86.6814 | 47.9987 |
| crRNA MFE | 72.5480 | 69.6068 |
| Unpaired probability | 72.1078 | 57.5635 |
| Local C<sub>min</sub> probability | 56.9153 | 37.2137 |
| hybridization MFE nt 15-23 | 54.5793 | 94.9857 |
| Guide G nt probability | 47.7324 | 29.2452 |
| Guide CA di-nt probability | 47.6241 | 21.4664 |
| hybridization MFE nt 3-15 | 47.3083 | 46.7462 |
| Guide CG di-nt probability | 44.0711 | 11.7069 |
| Guide AU di-nt probability | 42.1128 | 15.6205 |
| Guide CU di-nt probability | 40.6967 | 16.9172 |
| Guide A nt probability | 39.7297 | 25.0736 |
| Local GC<sub>max</sub> probability | 39.7236 | 59.0805 |
| Local AU<sub>min</sub> probability | 38.8318 | 56.3599 |
| Local U<sub>max</sub> probability | 38.7535 | 24.4657 |
| Guide GG di-nt probability | 38.5896 | 22.2621 |
| Local G<sub>min</sub> probability | 36.5244 | 19.6159 |
| Guide UC di-nt probability | 36.3043 | 15.2075 |
| Guide AC di-nt probability | 36.2901 | 14.2361 |
| Guide C nt probability | 36.0901 | 16.8637 |
| Local U<sub>min</sub> probability | 35.6886 | 16.2571 |
| Guide AG di-nt probability | 34.9534 | 14.6260 |
| Local A<sub>min</sub> probability | 34.6632 | 17.4928 |
| Local A<sub>max</sub> probability | 33.6557 | 16.1888 |
| Guide AA di-nt probability | 33.2106 | 13.3830 |
| Guide GC di-nt probability | 33.0670 | 13.3209 |
| Guide CC di-nt probability | 32.2834 | 12.9797 |
| Guide UG di-nt probability | 31.9188 | 13.1659 |
| Guide GU di-nt probability | 31.8413 | 12.8934 |
| Guide GA di-nt probability | 31.3837 | 13.1399 |
| Guide UA di-nt probability | 30.6930 | 12.5052 |
| Local C<sub>max</sub> probability | 29.8186 | 14.2189 |
| Local AU<sub>max</sub> probability | 22.4842 | 14.2585 |
| Predicted correctly folded DR | 8.2669 | 4.0679 |
| Predicted G-quadruplex | -3.0274 | 0.0219 |

### *Rfx*Cas13d guide scoring

We created a user-friendly R script that readily predicts *Rfx*Cas13d on-target guide scores. The only user-provided argument is a single-entry FASTA file input of minimally 30nt that represents the target sequence, such as a transcript isoform sequence. The software first generates all possible 23mer guide RNAs and collects all required features and predicts guide RNA efficacies. The only filter applied removes guide RNAs with homopolymers of 5 or more Ts and 6 or more Vs (V = A, C, G). Such guide RNAs may trigger early transcript termination for PolIII transcription or cause difficulties during oligo synthesis. The software returns a FASTA file with guide RNA sequences ranked by the predicted standardized guide score. In addition, a csv file is created following providing additional information. Optionally, the script can be used to plot the guide score distribution along the provided target sequence for visualization.

We used this software to predict guide scores for all transcripts (including all biotypes: `protein_coding, nonsense_mediated_decay, non_stop_decay, IG_*_gene, TR_*_gene, polymorphic_ pseudogene`) of protein coding genes annotated in GENCODE v19 (GRCh37) (*n* = 94,873 transcripts) and provide the top 10 ranked 5’UTR, coding sequence and 3’UTR annotating guide RNA sequences (**Supplementary Data 8**). We have made all guide score predictions available online (https://cas13design.nygenome.org).

### *Rfx*Cas13d guide scoring validation

To validate our that our initial RF_GFP_ model can readily separate between poorly and well-performing crRNAs, we performed several experiments.

First, we chose two genes that encode for cell surface proteins that allow for quantitative assessment of their expression levels by FACS. For each gene we predicted crRNAs for the highest expressed transcript isoform in HEK293FT cells (CD46: ENST00000367042.1, CD71 [TFRC]: ENST00000360110.4). For each gene, we selected 3 guides present in the low scoring quartiles (Q1 and Q2) and 3 guides in the high scoring quartiles (Q3 and Q4). We selected the guides to be non-overlapping and to reside in 3 different regions of the target transcript.

Then, we performed two essentiality screens with a dropout growth phenotype readout in HEK293FT and A375 cells, respectively. We designed two crRNA libraries targeting essential and control genes with a number of predicted low-scoring and high-scoring guide RNAs as described above (see **Screen library design and pooled oligo cloning**). For the HEK293FT cell screen, we compared the guide depletion of four groups of 30 guides (Essential gene targeted by high-scoring guide or by low-scoring guide, and control genes targeted by high-scoring guide or by low-scoring guide). We expected the greatest depletion for the 30 high-scoring guide RNAs targeting essential genes. Similarly, we compared the relative guide depletion of the same four groups of guide RNAs in the A375 screen, with the expectation that the 20 high-scoring guides per essential gene would be the most depleted.

For gene ranking based on guide depletion, we used robust rank aggregation (RRA) ^38^ to assign a *p*-value based on the consistency of log_2_FC-based rank-consistency of the most depleted *N* guide RNAs per gene (with *N* in {1, 5, 20}) across the two A375 screen replicates. The −log_10_ transformed *p*-values were then compared to other growth screens (RNAi and Cas9) using Spearman rank correlation. Specifically, we compared the RRA-derived log_10_ *p*-value to the log_2_FC from an RNAi-based DEMETER2 v5 repository ^27^ and the merged STARS scores from a Cas9-based approach^28^. For the correlation we only used genes with value present in all scores (Essential genes: *n* = 35; Control genes: *n* = 15).

Furthermore, we used the log_2_FC guide depletion values to compare the predictive value of the RF_GFP_ and RF_combined_ models. Specifically, for both essentiality screens we used 10 essential genes (all in HEK293FT and the 10 most depleted in A375 cells) and correlated the predicted guide scores from both models to the observed log_2_FC guide depletion scores (normalized to 0-100% per gene) of all detected guide RNAs (HEK293FT: *n* = 60 with 6 guides per gene; A375: *n* = 398 with up to 40 guides per gene). We made the same comparison on a per-gene level using all 40 guide RNAs per gene in the A375 screen.

### Data representation

In all boxplots, boxes indicate the median and interquartile ranges, with whiskers indicating either 1.5 times the interquartile range, or the most extreme data point outside the 1.5-fold interquartile. All transfection experiments show the mean of three replicate experiments with individual replicates plotted as points.

### Data availability statement

Screen data are being deposited to GEO with an accession number pending. Code to reproduce our analyses and figures are available on our gitlab repository (https://gitlab.com/sanjanalab/cas13). We also have provided pre-computed guide RNA predictions for guide RNAs targeting all protein-coding transcripts in the human genome on our web-based repository (https://cas13design.nygenome.org).

### Code availability statement

The predictive on-target model as well as all code for the analyses presented in the paper is available on our gitlab repository (https://gitlab.com/sanjanalab/cas13).

## Acknowledgements

We thank the entire Sanjana laboratory for support and advice. We thank R. Satija for helpful feedback, D. Knowles for discussions regarding predictive models, and M. Zaran for assistance with the web server. N.E.S. is supported by NYU and NYGC startup funds, NIH/NHGRI (R00HG008171, DP2HG010099), NIH/NCI (R01CA218668), DARPA (D18AP00053), the Sidney Kimmel Foundation, the Melanoma Research Alliance, and the Brain and Behavior Foundation. A.M.-M. is supported by CONACyT-Mexico Fellowship (#412653).

## Authors contributions

H.H.W and N.E.S conceived the project. H.H.W., N.E.S and A.M.-M. designed the experiments. A.M.-M. and H.H.W. performed the experiments. A.M.-M. and H.H.W analyzed experiments. H.H.W. analyzed the screen data, built the guide RNA prediction software and the online repository. X.G., M.L. and Z.D. helped with post-screen validation experiments. N.E.S. supervised the work. H.H.W. and N.E.S. wrote the manuscript with input from all of the authors.

