## Supplementary Figures and Notes for "Principles for rational Cas13d guide design"

**Supplementary Data**

| Supplementary Data | Description |
|---|---|
| 1 | Oligonucleotides |
| 2 | Guide and processed cells statistics |
| 3 | Screen read processing statistics |
| 4 | Raw guide RNA counts |
| 5 | Final guide RNA counts (after normalization; batch-correction; outlier-removal) |
| 6 | Guide RNA enrichments |
| 7 | Combined on-target model input including all features |
| 8 | Guide RNA predictions for protein coding transcripts in GENCODE v19 |

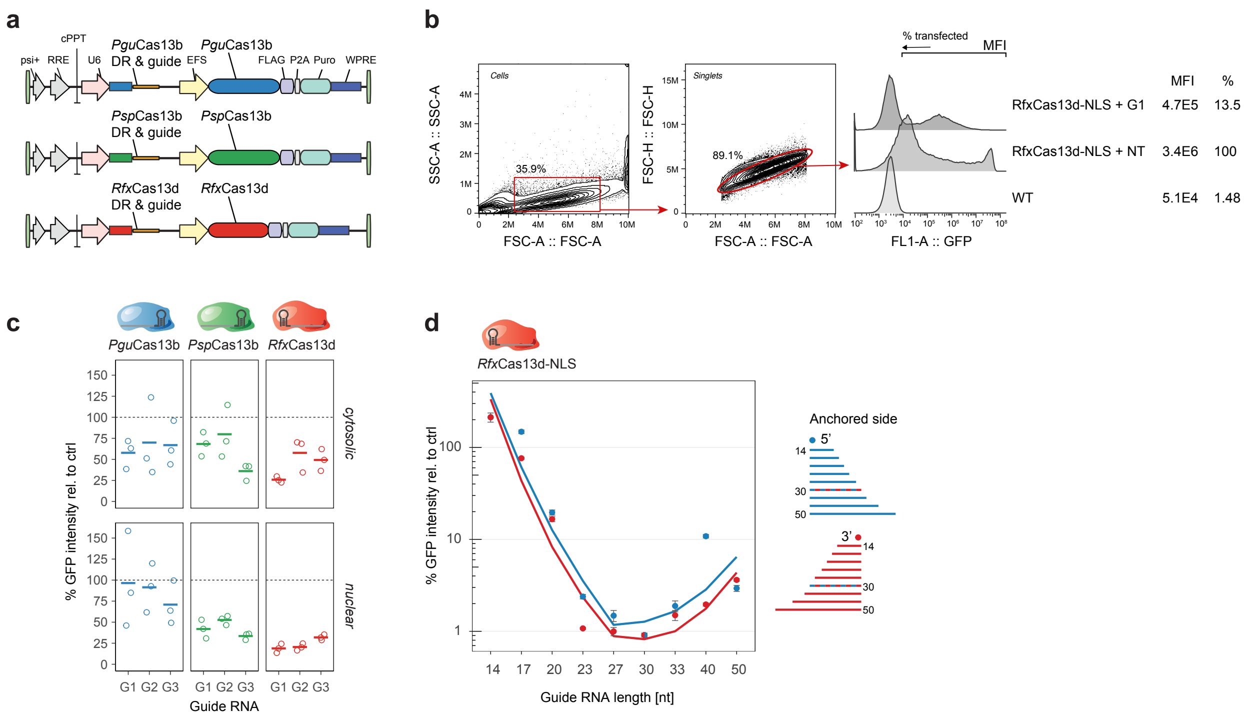


**Supplementary Figure 1. Comparison of Type IV CRISPR Cas protein knock-down efficacy.** (**a**) Lentiviral vectors for combined CRISPR Type IV enzyme and crRNA delivery. We generated single-vector effector protein plus crRNA expressing constructs utilizing nuclear localization (NLS) or cytosolic/nuclear-export signals (NES) to compare knock-down efficacies with uniform delivery, promoter and polyadenylation. (**b**) Gating strategy for transfection experiment presented in c. (**c**) Target knock-down comparison for *Pgu*Cas13b (blue), *Psp*Cas13b (green) and *Rfx*Cas13d (red) C-terminally fused to a NES (top) or NLS (bottom). Cells were co-transfected with plasmids encoding a Cas13 enzyme together with a crRNA, as shown in (*a*), and with a destabilized GFP plasmid. GFP intensity was recorded by fluorescence activated cell sorting 48 hours after transfection. Shown is the percentage of mean fluorescence intensity reduction of cells transfected with one of three different GFP-targeting guide RNAs sequences (G1, G2, G3) relative to a non-targeting guide RNA sequence for the same Cas13-fusion protein as a mean of three replicate experiments. Bars shown indicate the mean. (**d**) Target knock-down with different guide RNA lengths, while maintaining a fixed 5' or 3' anchor. *Rfx*Cas13d-NLS expressing HEK293 cells were co-transfected with plasmids delivering the crRNA only and a GFP expression plasmid. Shown is the percentage of mean fluorescence intensity reduction of cells transfected with a GFP-targeting guide relative to a non-targeting guide as a mean of three replicate experiments.

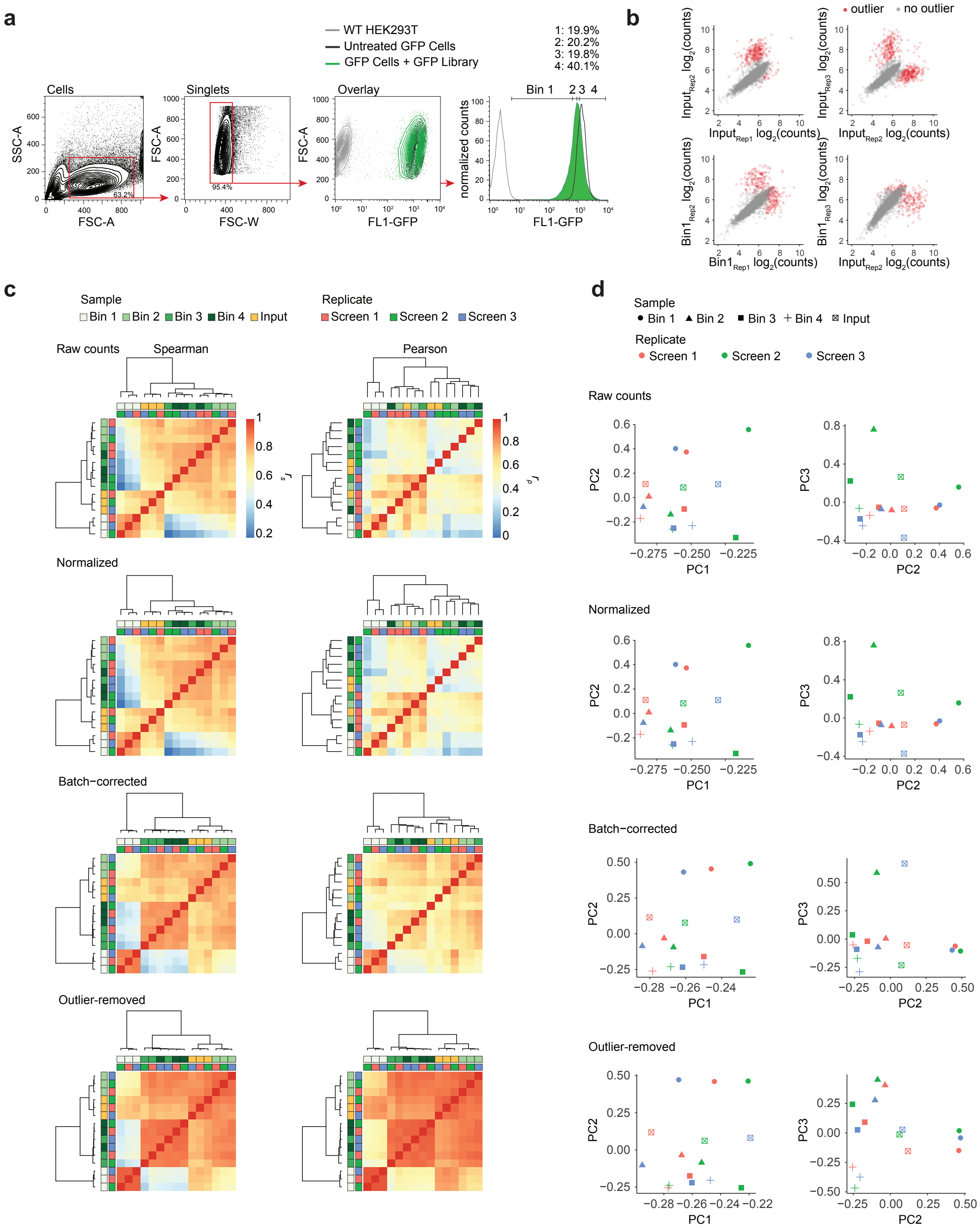


**Supplementary Figure 2. Processing of crRNA count data from the GFP pooled screen** (**a**) Gating strategy for GFP tiling screen (see methods). (**b**) Log2-transformed crRNA counts for three replicate screen input and bin 1 libraries showing flagged technical and non-reproducible outlier counts with high residuals deviating from a linear regression model. Flagged crRNAs counts of individual samples were not considered during crRNA count processing (see outlier removal in Supplementary Methods). (**c**) Spearman ($r_s$) and Pearson ($r_P$) correlations coefficients of crRNA counts across all 15 libraries during crRNA count processing. While Spearman correlations on the raw count data and terminally processed data looked comparable, outlier-removal greatly increase Pearson correlations between replicates and emphasized distance between different bins. (**d**) Principal components (PC) 1-3 corresponding to crRNA processing steps in (c).

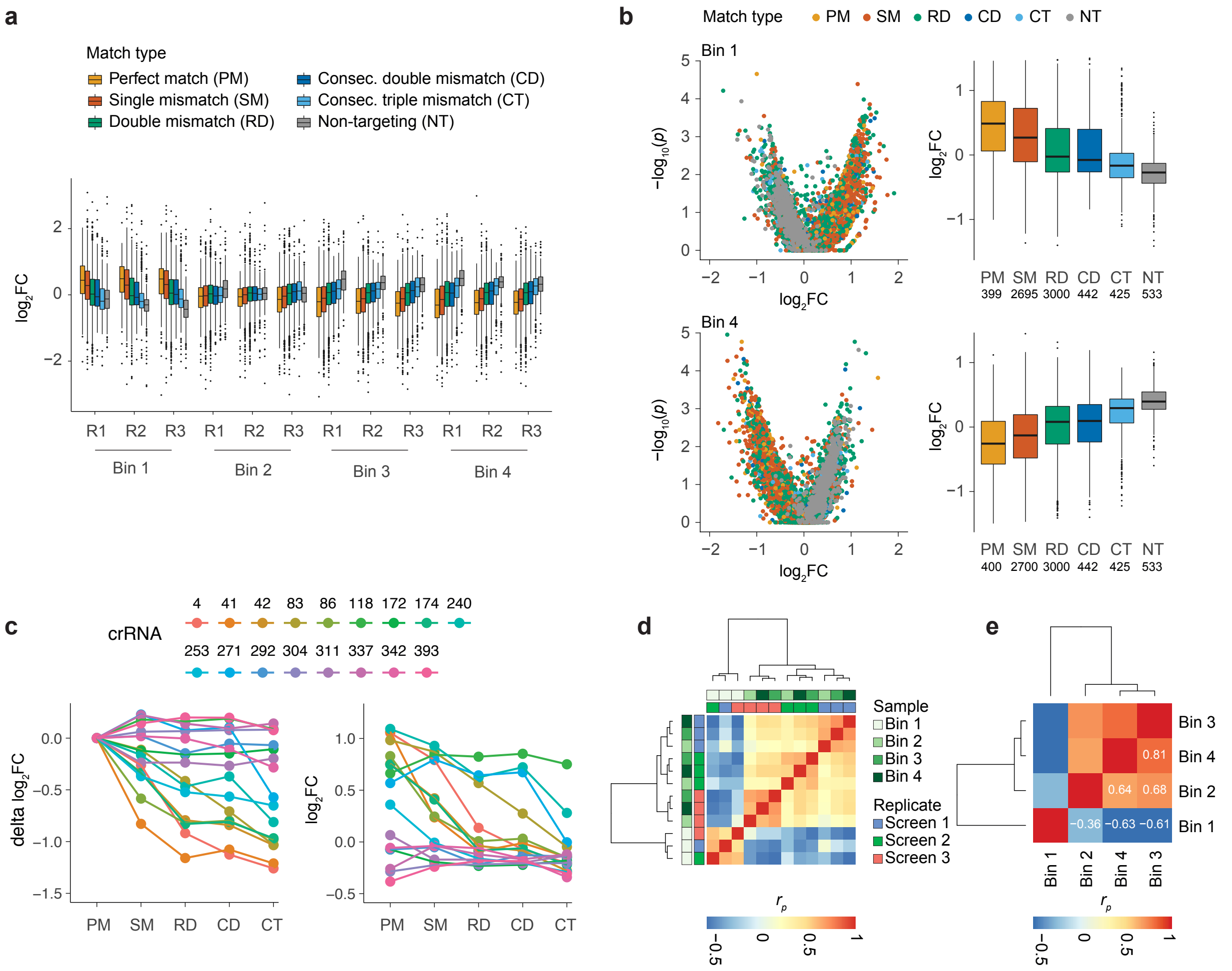


**Supplementary Figure 3. Guide RNA enrichments by guide type**. (**a**) $Log_2$ fold change ($log_2FC$) guide RNA count enrichment of each collected bin (Bin 1 – Bin 4) relative to the corresponding input sample. The guide RNA representation is segregated by guide RNA categories. (R = replicate screen). (**b**) $Log_2FC$ enrichment aggregated across all three replicates per bin. (*left*) Volcano plot showing $-log_{10}$-transformed $p$-values ($y$-axis) using robust rank aggregation (RRA) and the mean $log_2FC$ for each guide RNA separated by guide RNA match type. (*right*) Boxplot summarizing the mean $log_2FC$ by guide RNA match type ((*top*) Bin 1, (*bottom*) Bin 4). (**c**) (*left*) Delta $log_2FC$ of mismatch guide RNA minus its corresponding perfect match crRNA shown as a median for each guide RNA match type (PM $n = 1$, SM $n = 27$, RD $n = 100$, CD $n = 26$, CT $n = 25$). (*right*) As in *left*, but depicting the median $log_2FC$. (**d**) Pearson correlation coefficient ($r_p$) of $log_2FC$ enrichments for all bins as in (a). (**e**) As in (*d*), but using $r_p$ for the $log_2FC$ for each aggregated bin as in (*b*).

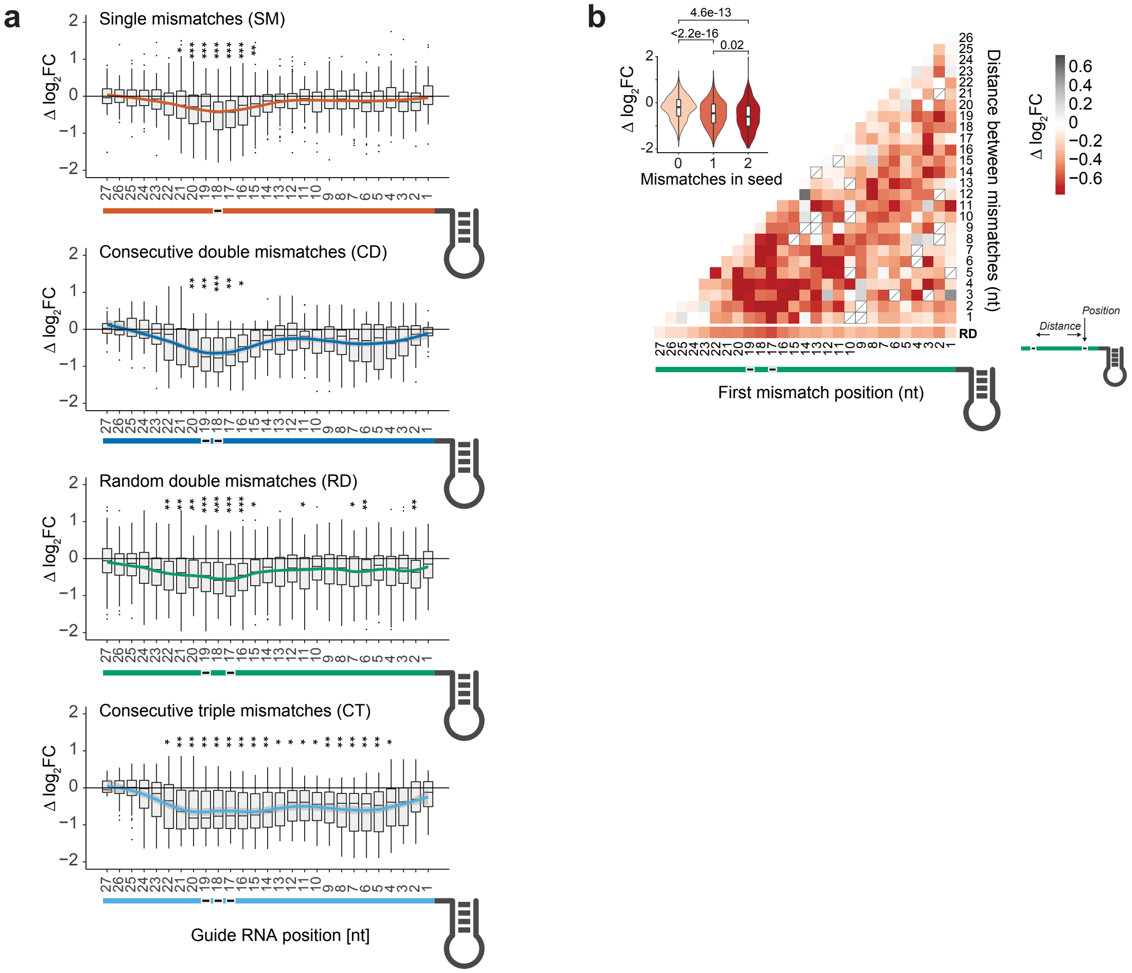


**Supplementary Figure 4. Identification of a critical guide RNA region for *Rfx*Cas13d guide RNAs.** (**a**) The decrease in targeting efficacy (delta $\log_2$FC) of crRNAs with single nucleotide mismatches, consecutive double mismatches, random double mismatches and consecutive triple mismatches relative to their cognate perfectly matching guides stratified by mismatch position (two-sided *t*-test of $\log_2$FC values of permutated guide RNAs versus perfect match guide RNAs. Significance levels: * $p < 0.05$, ** $p < 0.01$, *** $p < 0.001$). (**b**) Delta $\log_2$FC in targeting efficacy for random double (RD) mismatches. RD mismatched guides are plotted by the position of the first mismatched base (*x-axis*) and the distance between the first and second mismatched bases (*y-axis*). The inset shows the relative change of targeting efficacy summarized by the number of mismatches falling into the seed region (positions 15 to 21).

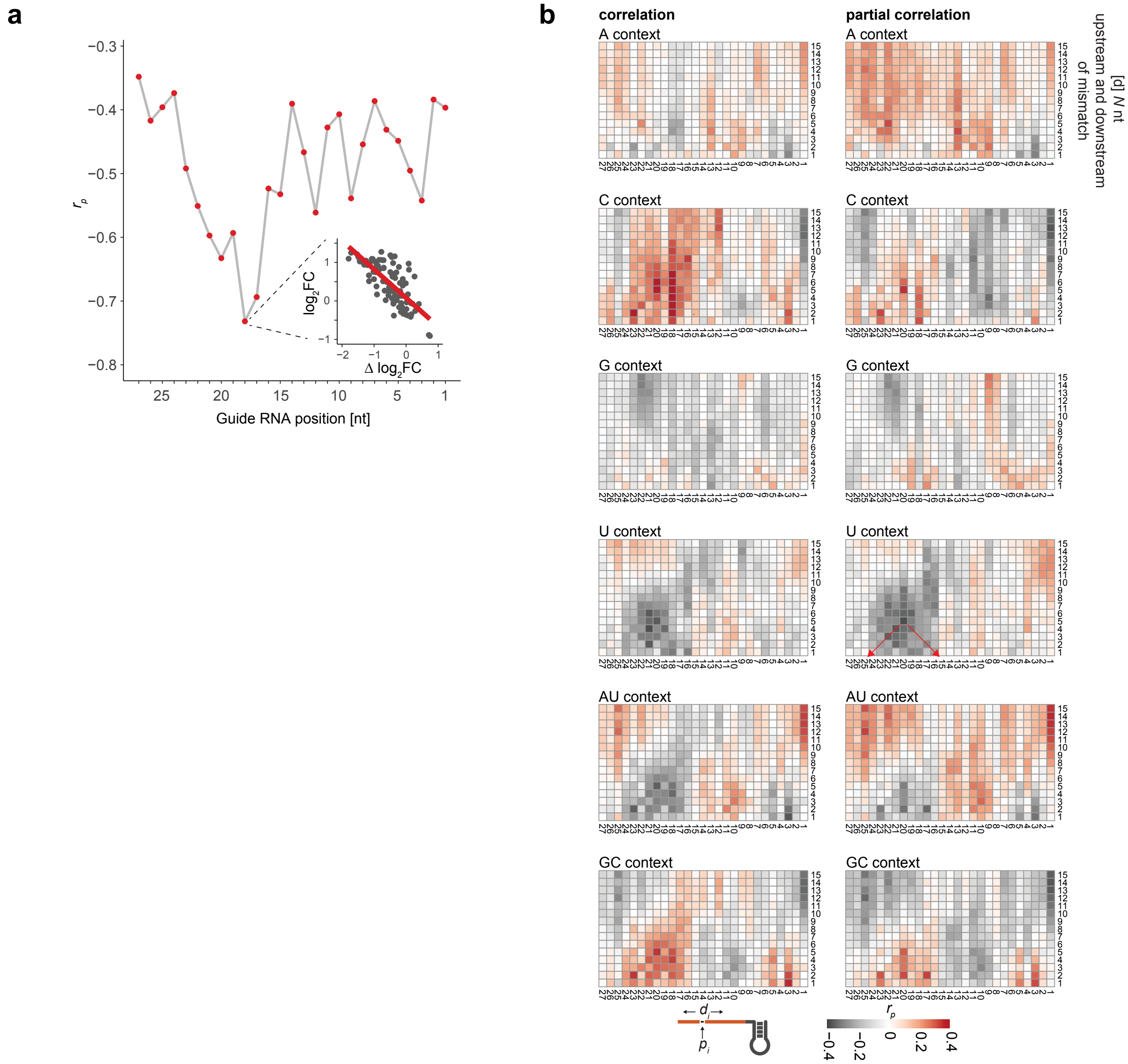


**Supplementary Figure 5. The nucleotide context around single nucleotide mismatches influences the mismatch tolerance.** (**a**) Pearson correlation coefficient ($r_p$) between observed $log_2FC$ and delta $log_2FC$ for all single mismatch guide RNAs relative to their cognate perfect matching guide RNAs segregated by all 27 positions. (*inset*) Scatterplot depicting the relationship between observed $log_2FC$ and delta $log_2FC$ for all guide RNAs with a mismatch at position 18 relative to the 5’ guide RNA end. (**b**) Heatmaps showing the correlation (*left*) and partial correlation controlling for crRNA folding minimum free energy (MFE) (*right*) of the observed crRNA $log_2FC$ and the nucleotide context (A, C, G, U, A|U, G|C) around single nucleotide mismatches. For each mismatch position $p$ relative to the 5’ guide RNA end the nucleotide density was calculated as a fraction in a window extended by $d$ (1 - 15 nt) on both sides centered on the mismatch position $p$.

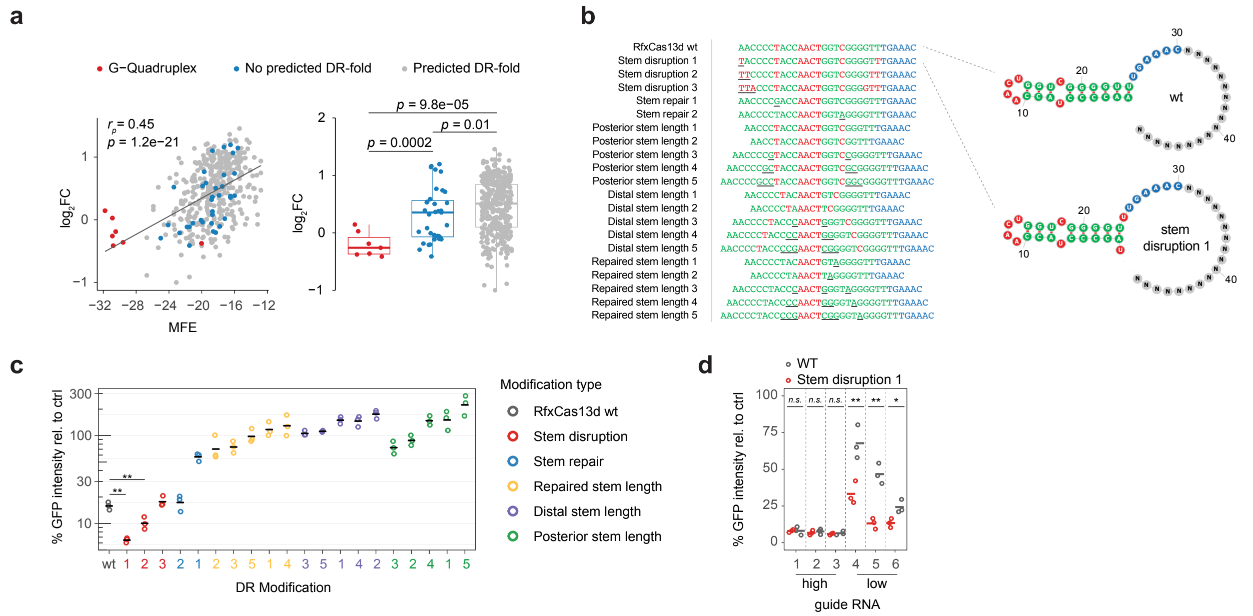


**Supplementary Figure 6. crRNA structure affects crRNA targeting efficacy.** (**a**) Scatterplot showing the crRNA $\log_2$FC versus the predicted crRNA secondary structure minimum free energy (MFE). The Pearson correlation coefficient ($r_p$) is nearly unchanged ($r_p$ = 0.44) when MFEs of G-Quadruplex-forming crRNAs are ignored. (**b**) (left) DR modification tested in (c). Key: green = paired nucleotide, red = unpaired nucleotide in bulge or loop, blue = unpaired nucleotide preceding guide RNA, underlined bases have been changed relative to the wild-type reference. Deleted bases are not shown. (right) Predicted DR secondary structure of two selected DR modifications using RNAfold. (**c**) Target knock-down comparison varying the DR sequence using GFP-targeting guide G3 used in **Supplementary Fig. 1c**. *Rfx*Cas13d-NLS expressing cells were co-transfected with plasmids delivering the crRNA and with a GFP-encoding plasmid. Shown is the percentage of mean fluorescence intensity reduction of cells transfected with a GFP-targeting guide relative to a non-targeting guide as a mean of three replicate experiments. Error bars indicate standard error of the mean. (**d**) Target knock-down comparison comparing the wildtype DR (WT) and stem disruption 1 DR (as in b) across 6 GFP-targeting guide RNAs with either low or high knock-down relative to a non-targeting guide control (Guides as used in **Fig. 1e**). ($n$ = 3 transfection replicates). One-tailed t-test in *c* and *d* with significance levels: * $p < 0.05$, ** $p < 0.01$, *** $p < 0.001$)

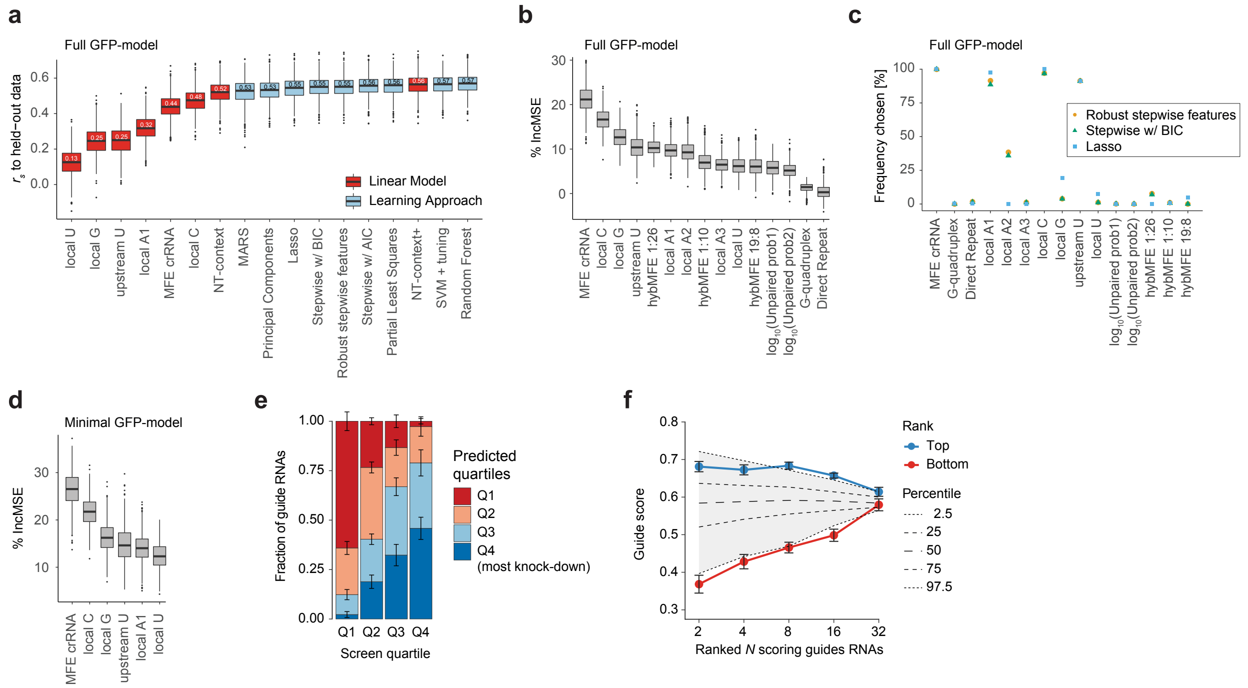


**Supplementary Figure 7. Comparison of machine learning approaches and feature selection for Cas13d guide RNA 'on-target' model.** (**a**) Model performance evaluation of linear models and learning approaches using features as in **Supplementary Table 1**. We compared the ability of machine leaning regression approaches to predict target knock-down of held-out data using bootstrapping. The data of all perfectly matching guides (n = 399) from the GFP tiling screen and features was randomly split into 70% training data and 30% held-out testing data for 1000 random non-redundant splits. The prediction accuracy (comparing predicted scores to the known $log_2FC$) is computed using the Spearman correlation ($r_s$) to the held-out data. The number inside the box is the median of 1000 bootstrap samplings. The models are ranked by their median performance. (NT-context = linear combination of all nucleotide context values. NT-context+ = NT-context plus crRNA MFE). (**b**) Boxplots showing the percent increase in mean-squared error (%IncMSE) of features for the top-scoring Random Forest model in (a). (**c**) Feature selection of selected machine learning approaches. Shown is the frequency that a particular feature was selected for the best model across 1000 iterations from (a). (**d**) %IncMSE of features for the top-scoring Random Forest model using a minimal set of selected features, corresponding to the $RF_{minimal}$ (=$RF_{GFP}$) model in **Fig. 2a**. (**e**) Comparison of predicted and measured fold-change quartiles across the 10-fold $RF_{GFP}$ model cross-validation. (**f**). Prediction of standardized guide scores by the $RF_{GFP}$ model. Shown are the median predicted guides scores for the top/bottom guide RNAs (N = 2, 4, 8, 16, 32) ranked by the known guide RNA efficacy across the 10-fold cross-validation. Error bars indicate the standard error of mean. Grey shading indicates the null distribution for the median guides scores of randomly selected crRNAs across 1000 samplings for each N.

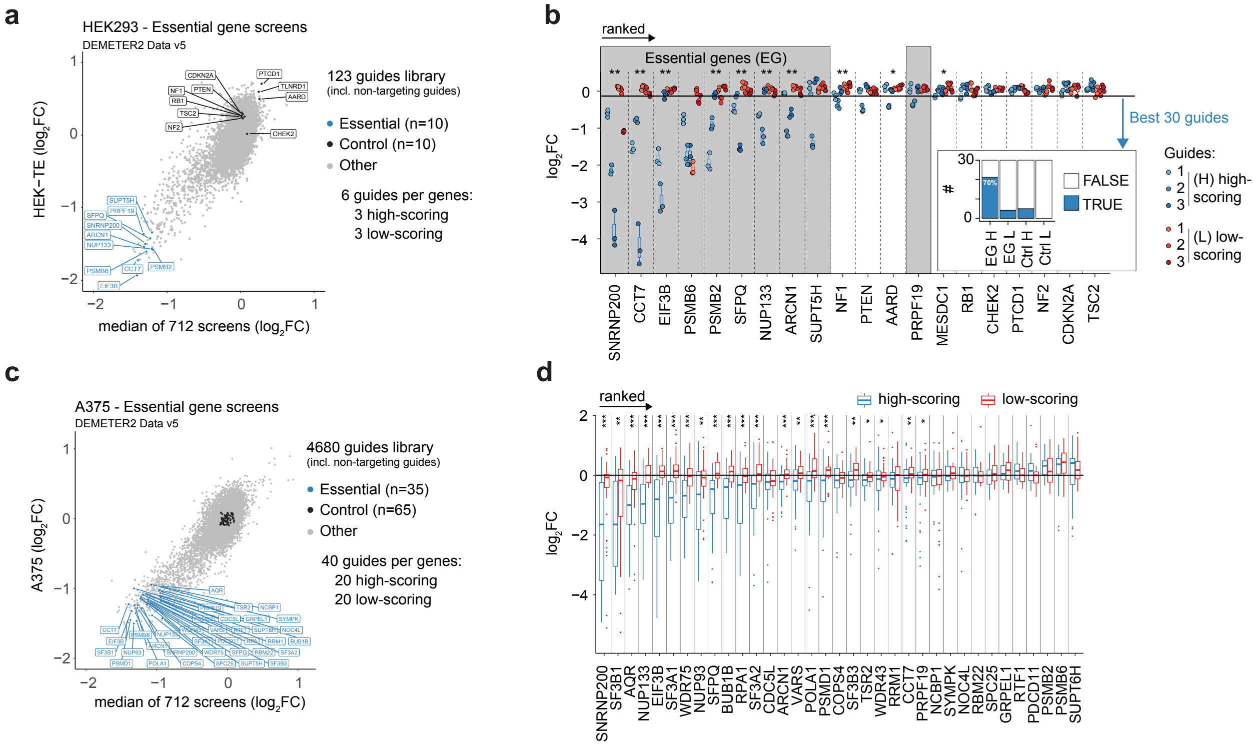


**Supplementary Figure 8. RF$_{GFP}$ on-target model validation.** (**a**). Scatterplot showing the $\log_2$FC depletion scores provided by the DEMETER2 v5 repository of HEK-TE cells (y-axis) and the median $\log_2$FC across 712 cell lines present (x-axis). Genes targeted in the HEK293 cell essentiality screen are colored by gene type (essential, control). (**b**) Guide RNA depletion ($\log_2$FC) in pooled HEK293 cell essentiality screen at Day 27 relative to input samples. Shown are the individual guide enrichments colored predicted guide scoring class (high vs. low) of three replicate experiments. Genes are ranked by the mean depletion of the predicted high-scoring guides. (inset) Distribution of the 30 most depleted guide RNAs across four targeting classes (EG H: High-scoring guide RNAs targeting essential genes; EG L: Low-scoring guide RNAs targeting essential genes; Ctrl H: High-scoring guide RNAs targeting non-essential genes; Ctrl L: Low-scoring guide RNAs targeting non-essential genes). (**c**) Selected target genes for A375 cell essentiality screen similar to a. *(***d***)*. Boxplot summarizing the $\log_2$FC depletion of predicted low- and high-scoring guide RNAs for the 35 selected essential genes comparing Day 14 to input samples. For *b* and *d* we used a one-sided t-test of $\log_2$FC values of high-scoring guide RNAs versus low-scoring guide RNAs per gene. Significance levels: * $p < 0.05$, ** $p < 0.01$, *** $p < 0.001$.

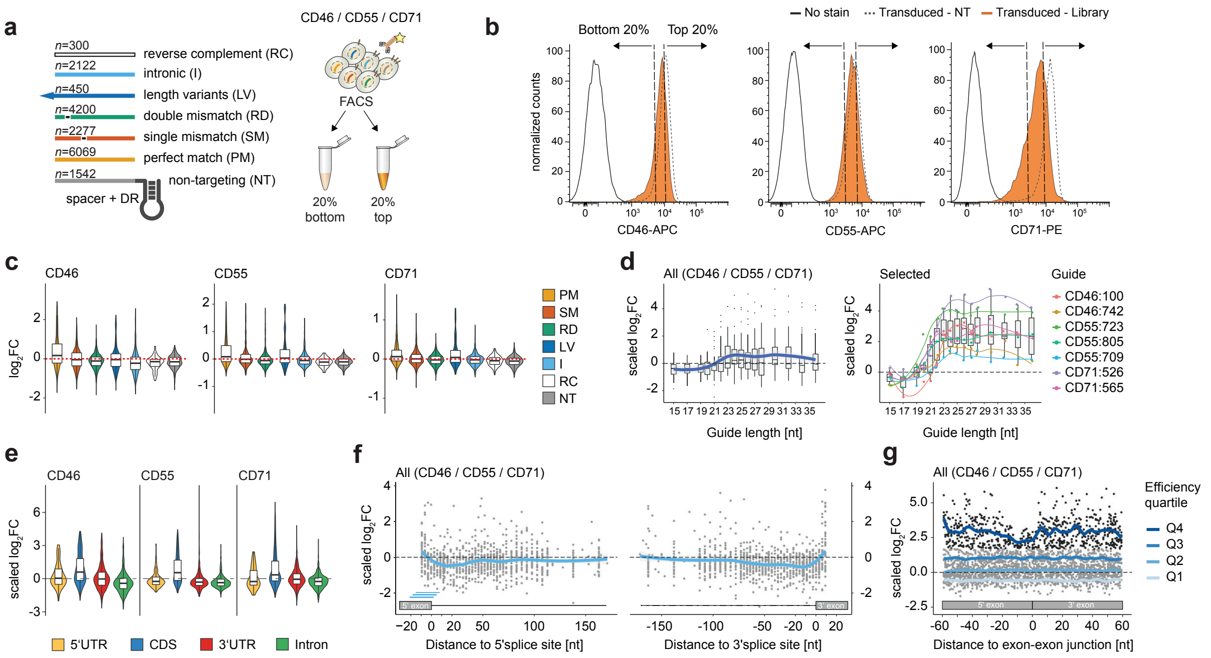

**Supplementary Figure 9. *Rfx*Cas13d cell surface protein knock-down screens.** (**a**) Together, the three libraries (CD46, CD55, CD71) contained 6,069 guide RNAs that were perfectly-matched, 99 guide RNAs with a single mismatch at each of the 23 guide positions ($n$ = 2,277 guides), 42 guide RNAs with 100 random double mismatches (n = 4,200 guides), and 30 guides with 10 additional guide length variants (n = 450 guides), a set of 2188 intron targeting guides centered around the splice-donor and acceptor sites across 40 introns and 300 reverse complement perfect match guide RNAs as additional negative control. For details, see **Supplementary Data 2**. CrRNAs are lentivirally transduced into TetO-*Rfx*Cas13d HEK293 cells. Five to ten days after transduction, cells are stained for the targeted cell surface protein and sorted by intensities into 2 bins (Bottom 20% = strongest knock-down, Top 20% = least knock-down). (**b**) Sorting strategy applied for cell surface protein tiling screens comparing unstained cells, cells transduced with the respective crRNA library or with a single non-targeting guide RNA. (**c**) $Log_2$ fold-change ($log_2FC$) enrichment scores of guide RNAs comparing guide RNA counts of the bottom 20% cell surface protein intensity bin (Bin 1) to the input (unsorted) cell population. Scores are demarcated by the type of designed guide RNAs as given by the list in *a*. (**d**) $Log_2FC$ guide enrichments of 3'end guide length variants grouped by guide RNA length. (*left*) All length variants highlighting the mean behavior using LOESS regression. (*right*) Selected guide RNAs. (**e**) $Log_2FC$ enrichments of perfect match guide RNAs grouped by annotation category for each cell surface protein. (**f**) $Log_2FC$ guide enrichments of intron-targeting guide RNAs across all 40 introns present in the three target genes. Guide RNAs were densely spaced around the 5' splice-site (*left*) or 3'splice-site (*right*) with a minimum of 1 nucleotide falling into the intron and up to 150 nucleotide distance to the exon. Values are depicted relative to the midpoint of the guide RNA. The mean behavior is highlighted using LOESS regression. (**g**) $Log_2FC$ guide enrichments of guide RNAs with target-sites close to exon-exon junctions across all 40 junctions present in the three target genes. Values are depicted relative to the midpoint of the guide RNA. The mean behavior is highlighted using LOESS regression separated into quartiles to highlight positional depletion of strongly enriched guide RNAs directly upstream to the junction specific to the highest quartile Q4. For *d-g*, $log_2FC$ values were scaled within each screen.

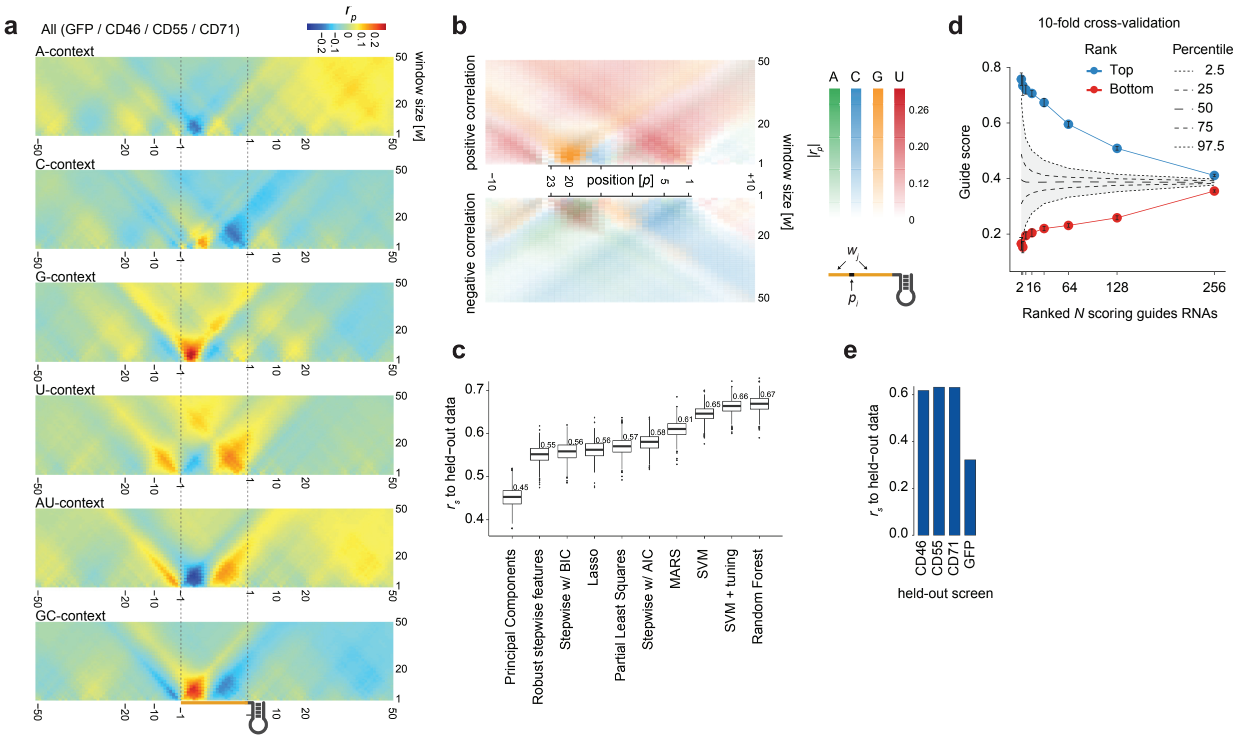


**Supplementary Figure 10. Nucleotide preferences and combined on-target model performance. (a)** Heatmaps depicting the Pearson correlation coefficient ($r_p$) between the local target nucleotide-contexts (A, C, G, U, A|U and G|C) and observed $log_2FC$ relative to guide RNA match positions. We performed a grid-search correlating the observed guide RNA efficacies with the target nucleotide probabilities across a window of 1 nt up to 50 nt at every point 75 nt upstream to 75 nt downstream relative to all 2,918 selected CDS-targeting perfect match target sites. See inset diagram in *b* for schematic of the window. Color scale applies to all six heatmaps. We determined the for each nucleotide (A,C,G,U, A|U, G|C) the position and widow size of minimal and maximal Pearson correlation coefficient (**Supplementary Table 2**). Patterns derived from partial correlation controlling for the crRNA MFE did not deviate from correlations shown. (**b**) Similar to *a* overlaying the positive (*top*) and negative (*bottom*) correlation coefficients in a transparent-to-color scale separated by nucleotide to highlight mutually exclusive position specific correlations. (**c**) Comparison of machine leaning regression approaches to predict target knock-down of held-out data using bootstrapping. The data of all CDS-targeting perfect match guides ($n$ = 2,918) from the all four tiling screens and features (**Supplementary Table 2**) was randomly split into 70% training data and 30% held-out testing data for 1000 random non-redundant splits. The prediction accuracy (comparing predicted scores to the known $log_2FC$) is computed using the Spearman correlation ($r_s$) to the held-out data. Models are ranked by their median performance. (**d**) Prediction of standardized guide scores by the $RF_{combined}$ model. Shown are the mean predicted guides scores for the top/bottom guide RNAs ranked by the known guide RNA efficacy across the 10-fold cross-validation. Error bars indicate the standard error of mean. Grey shading indicates the null distribution for the mean guides scores of randomly selected guide RNAs across 1000 samplings. (**e**) Spearman rank correlation between measured guide efficiency ($log_2FC$) and the predicted guide score for the indicated gene using leave-one-out cross-validation.

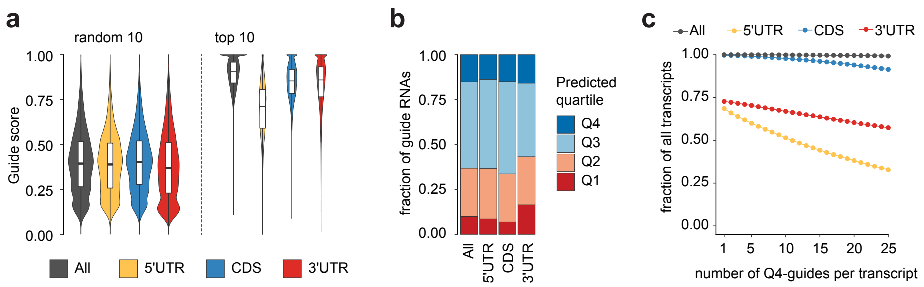


**Supplementary Figure 11. Transcriptome-wide *Rfx*Cas13d crRNA guide prediction.** (**a**) Predicted score distributions of 10 randomly sampled (*left*) and the top 10 guide RNAs (*right*) per transcript, overall and for each mRNA sub-annotation category. The top 10 guide RNAs per transcript are provided in **Supplementary Data 8**. The full set of predictions are available on https://cas13design.nygenome.org. (**b**) The predicted guide score quartiles according to the efficiency quartiles of the four combined tiling screens shown for 10,000 randomly sampled transcripts of protein-coding genes in GENCODE v19. (**c**) Fraction of processed transcripts that contain at least 1 (up to 25) prediction in the highest scoring quartile (Q4). (All = whole transcript).

**Supplementary Table 1. Input features for GFP 'on-target' model selection.**

| | Feature Name | Description |
|---|---|---|
| **1** | crRNA MFE | Minimum free energy value of DR-sequence plus guide sequence using `RNAfold` |
| **2** | direct repeat | Binary - based on the presence of the predicted DR fold "(((((.(((....))).)))))" at the crRNA start |
| **3** | G-quadruplex | Binary - based on the presence of the predicted G-quadruplex indicated by "+" within the folding sequence |
| **4** | hybMFE 1:26 | Minimum free energy value between guide RNA nucleotides 1-26 and its corresponding target site (=overall hybridization) |
| **5** | hybMFE 1:10 | Minimum free energy value between guide RNA nucleotides 1-10 and its corresponding target site (=5' hybridization) |
| **6** | hybMFE 19:8 | Minimum free energy value between guide RNA nucleotides 19-27 and its corresponding target site (=3' hybridization) |
| **7** | $\log_{10}$(Unpaired prob1) | $\log_{10}$(probability) of a target RNA nucleotide being unpaired in a window centered at nt -23 relative to the guide match start summarizing 21 nts (nt -13:-33) |
| **8** | $\log_{10}$(Unpaired prob2) | $\log_{10}$(probability) of a target RNA nucleotide being unpaired in a window centered at nt -23 relative to the guide match start summarizing 10 nts (nt -27:-18) |
| **9** | A1 context | Probability of target RNA A-bases at position -22 relative to the guide match start summarizing 7 nts (nt -19:-25) |
| **10** | A2 context | Probability of target RNA A-bases at position -22 relative to the guide match start summarizing 33 nts (nt -6:-48) |
| **11** | A3 context | Probability of target RNA A-bases at position -16 relative to the guide match start summarizing 20 nts (nt -25:-6) |
| **12** | C context | Probability of target RNA C-bases at position -11 relative to the guide match start summarizing 22 nts (nt -21:0) |
| **13** | G context | Probability of target RNA G-bases at position -10 relative to the guide match start summarizing 21 nts (nt 20:0) |
| **14** | U context | Probability of target RNA U-bases at position -3 relative to the guide match start summarizing 18 nts (nt -12:+5) |
| **15** | upstream U context | Probability of target RNA U-bases at position -78 relative to the guide match start summarizing 30 nts (nt -93:-64) |

For the $RF_{GFP}$ model we define features as follows: For guide RNA features (features **4**, **5**, **6**), nucleotide 1 defines the guide match start site (GMSS) being the most 5' guide RNA base matching the target RNA. Nucleotide 2 relative to GMSS is the subsequent base (moving in the 5' to 3' direction) in the guide RNA and so on. For target RNA features (features **7** – **15**), we denote the target nucleotide opposite to the GMSS as nucleotide 0. Moving in 5' to 3' direction target RNA nucleotide -1 is upstream (5') to target RNA nucleotide 0 and base-paired to guide nucleotide 2, while target RNA nucleotide +1 is downstream of the target site and so on. A complete illustration for features **4** – **15** with a schematic of the guide RNA and target RNA can be found in Supplementary Note 1, Figure 6.

**Supplementary Table 2. Selected Input features for $RF_{combined}$ 'on-target' model.**

| | Feature Name | Description |
|---|---|---|
| **1** | crRNA MFE | Minimum free energy value of DR-sequence plus guide sequence using `RNAfold` |
| **2** | Direct repeat | Binary - based on the presence of the predicted DR fold "(((((.(((....))).)))))" at the crRNA start |
| **3** | G-quadruplex | Binary - based on the presence of the predicted G-quadruplex indicated by "+" within the folding sequence |
| **4** | HybMFE 3:15 | Minimum free energy value between guide RNA nucleotides 3-15 and its corresponding target site (=5' hybridization) |
| **5** | HybMFE 15:23 | Minimum free energy value between guide RNA nucleotides 15-23 and its corresponding target site (=3' hybridization) |
| **6** | $Log_{10}$(Unpaired prob) | $log_{10}$(probability) of a target RNA nucleotide being unpaired in a window centered at nt -11 relative to the guide match start summarizing 23 nts (nt 0:-22) |
| **7** | Local $A_{max}$ probability | Probability of target RNA A-bases at position -10 relative to the guide match start summarizing 8 nts (nt -14:-7) |
| **8** | Local $C_{max}$ probability | Probability of target RNA C-bases at position -16 relative to the guide match start summarizing 4 nts (nt -18:-15) |
| **9** | Local $G_{max}$ probability | Probability of target RNA G-bases at position -19 relative to the guide match start summarizing 3 nts (nt -20:-18) |
| **10** | Local $U_{max}$ probability | Probability of target RNA U-bases at position -6 relative to the guide match start summarizing 12 nts (nt -12:-1) |
| **11** | Local $AU_{max}$ probability | Probability of target RNA A or U-bases at position -7 relative to the guide match start summarizing 11 nts (nt -12:-2) |
| **12** | Local $GC_{max}$ probability | Probability of target RNA G or C-bases at position -18 relative to the guide match start summarizing 9 nts (nt -22:14) |
| **13** | Local $A_{min}$ probability | Probability of target RNA A-bases at position -17 relative to the guide match start summarizing 7 nts (nt -20:-14) |
| **14** | Local $C_{min}$ probability | Probability of target RNA C-bases at position -3 relative to the guide match start summarizing 9 nts (nt -7:+1) |
| **15** | Local $G_{min}$ probability | Probability of target RNA G-bases at position -9 relative to the guide match start summarizing 9 nts (nt -13:-5) |
| **16** | Local $U_{min}$ probability | Probability of target RNA U-bases at position -17 relative to the guide match start summarizing 10 nts (nt -22:-13) |
| **17** | Local $AU_{min}$ probability | Probability of target RNA A or U-bases at position -17 relative to the guide match start summarizing 9 nts (nt -21:-13) |
| **18** | Local $GC_{min}$ probability | Probability of target RNA G or C-bases at position -18 relative to the guide match start summarizing 11 nts (nt -11:-1) ($GC_{min}$ - *not used in $RF_{combined}$*) |
| **19-22** | Nucleotide probability | Probability of guide RNA A,C, G or U bases (U - *not used in $RF_{combined}$*) |
| **23-38** | Di-nucleotide probability | Probability of 16 possible guide RNA di-nucleotides (UU - *not used in $RF_{combined}$*) |

For the $RF_{combined}$ model we define features as follows: For guide RNA features (features **4** and **5**), nucleotide 1 defines the guide match start site (GMSS) being the most 5' guide RNA base matching the target RNA. Nucleotide 2 relative to GMSS is the subsequent base (moving in the 5' to 3' direction) in the guide RNA and so on. For target RNA features (features **6** – **18**), we denote the target nucleotide opposite to the GMSS as nucleotide 0. Moving in 5' to 3' direction target RNA nucleotide -1 is upstream (5') to target RNA nucleotide 0 and base-paired to guide nucleotide 2, while target RNA nucleotide +1 is downstream of the target site and so on. A complete illustration for features **4** – **18** with a schematic of the guide RNA and target RNA can be found in Supplementary Note 2, Figure 12.

### Supplementary Note 1

#### Features of Cas13d targeting from the GFP tiling screen

#### Anti-Tag

Recently, others have found that Cas13a is inhibited by a 4 nt "anti-tag" sequence — homology between the end of the DR and the corresponding flanking sequence of the target — and have speculated that Cas13d, which has a similarly positioned 5' DR, might also use an anti-tag for host versus pathogen discrimination [1]. Using all perfect match guide RNAs, we did not find evidence for the presence of a similar anti-tag sequence for *Rfx*Cas13d suggesting that anti-tags may not be found in all Type VI CRISPRs or contribute only marginally compared to other features (**Note Fig. 1**).

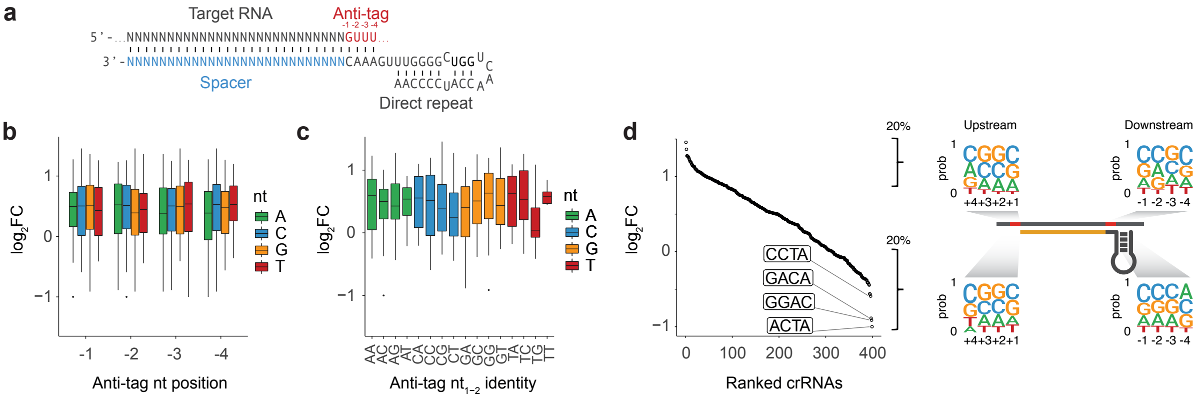


**Note Figure 1. No evidence for the presence of an *Rfx*Cas13d anti-tag in GFP tiling screen.** (**a**) Anti-tag position in target RNA. (**b**) $\log_2$FC of all perfect match guides ($n$ = 399) segregated by anti-tag nucleotide identity at anti-tag positions 1 through 4. (**c**) Same as in (*b*), but segregated by anti-tag di-nucleotide identity at anti-tag positions 1 and 2 (**d**) (*left*) $\log_2$FC-ranked guide RNAs highlighting the anti-tag sequence for the 4 least efficient guides. (*right*) Position Weight Matrices (PWMs) depicting the positional nucleotide probabilities of either the top 20% or bottom 20% $\log_2$FC-ranked guides 5' and 3' (i.e. putative anti-tag) relative to the guide RNA match position.

#### Nucleotide preferences

Next, we tested whether position-based nucleotide preferences exist within the guide RNA target sequence or nearby nucleotides by comparing the nucleotide composition of the top 20% to all perfectly matching guides in the GFP screen, similar to previous approaches assessing Cas9 guide preferences [2]. Although the top enriched guides showed slight nucleotide preferences compared to all guides, most preferences became insignificant after correction for multiple hypothesis testing (**Note Fig. 2a**). However, when we correlated guide RNA nucleotide probabilities with the observed guide enrichment, we saw that high 'G'-content in the guide RNA had a strong negative impact (**Note Fig. 2b-c**). Other measures, like guide RNA GC-content indicated a local optimum around 50% with lower guide efficiency when the guide adopts lower or higher GC proportions.

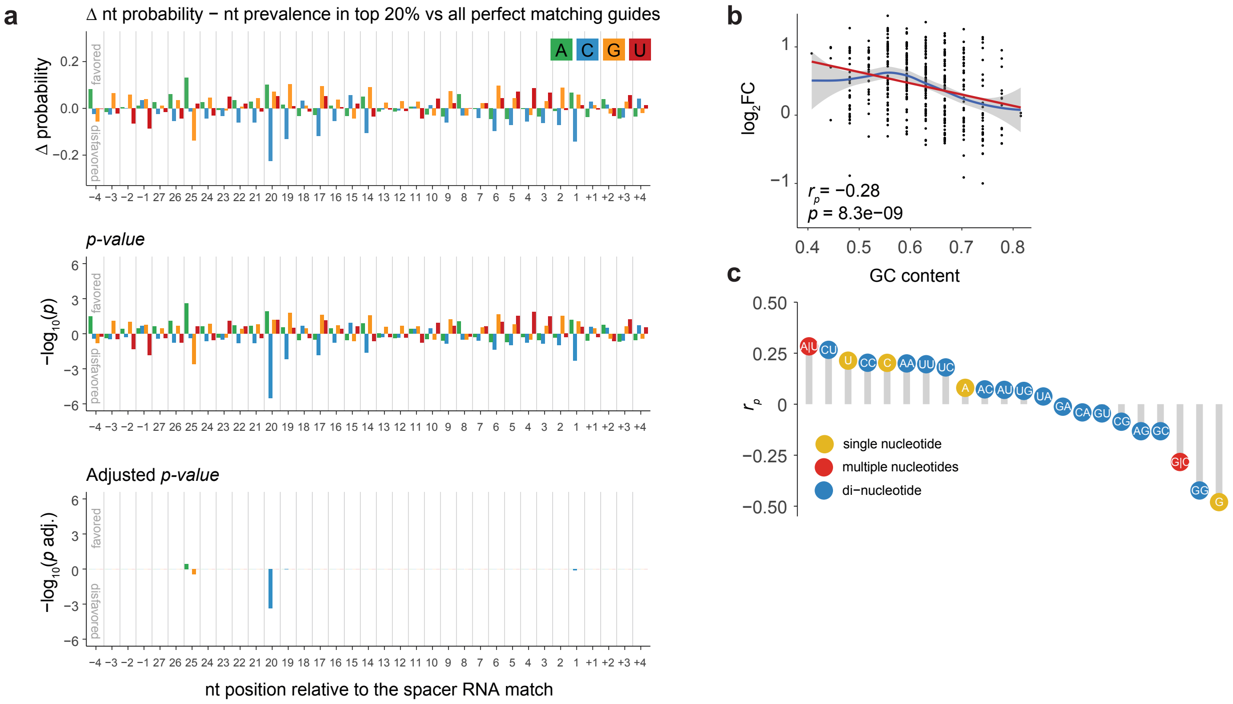


**Note Figure 2. Cas13d guide RNA nucleotide preferences influence guide RNA efficacies in GFP screen.** (**a**) (*top*) Effect-size (delta nucleotide probabilities), (*middle*) *p*-values and (*bottom*) Bonferroni-corrected *p*-values of observing the conditional probability of a guide in the top 20% under the null distribution examined at every position including the 4 nucleotides 5' and 3' of the guide RNA target site. The *p*-values were calculated from the binomial distribution with a baseline probability estimated from the full-length mRNA target sequence with all perfect match guide RNAs. (**b**) Scatterplot depicting the guide RNA $\log_2$FC and guide RNA GC-content as a fraction. The red line indicates the linear relationship between both values. The blue line indicates a LOESS fit and has a local optimum around a GC-content of 0.5-0.6. (**c**) As in *b*, but showing the Pearson correlation coefficient ($r_p$) between guide RNA $\log_2$FC and all guide single nucleotides, di-nucleotides, and G|C and A|U-content.

### crRNA folding

The negative correlation to the observed guide RNA enrichments ($\log_2$FC) was restricted to high 'G'-content in the guide RNA, while guide RNA 'C'-content did not affect targeting in the same way (**Note Fig. 2c**). This suggests that the effect may not be caused by specific guide-target interaction, which should weight 'C' and 'G' bases interchangeably, but instead may be driven by 'G'-dependent stable structures within the crRNA that may render the crRNA inaccessible for Cas13d. Indeed, predicting the secondary structure and corresponding minimum free energy (MFE) of perfect match guides showed a positive correlation between the MFE and guide efficacy (**Supplementary Fig. 6a**). In particular, 'G'-dependent structures, such as predicted G-quadruplexes, showed diminished target knock-down.

### Guide RNA – target RNA hybridization

We next tested whether guide-target hybridization can contribute to guide RNA efficacy by computing the correlation between hybridization energy and guide RNA efficacy (**Note Fig. 3a**). For the GFP screen we found that more stable hybridization between guide RNAs and their target sequences (lower MFE) was correlated with lower guide RNA efficacy ($r$ = 0.31) (**Note Fig. 3b**).

This suggests that the most stable guide-target interactions may render the ribonucleoprotein complex less active. Interestingly, calculating MFEs between smaller regions within the guide RNA indicated multiple sub-structures that contribute to the overall correlation, suggesting that individual parts of the guide-target interaction may serve specific roles during ternary complex formation or nuclease activation (**Note Fig. 3a**). However, these correlative structures were nearly gone when using partial correlation to control for the effect of crRNA folding (**Note Fig. 3c**).

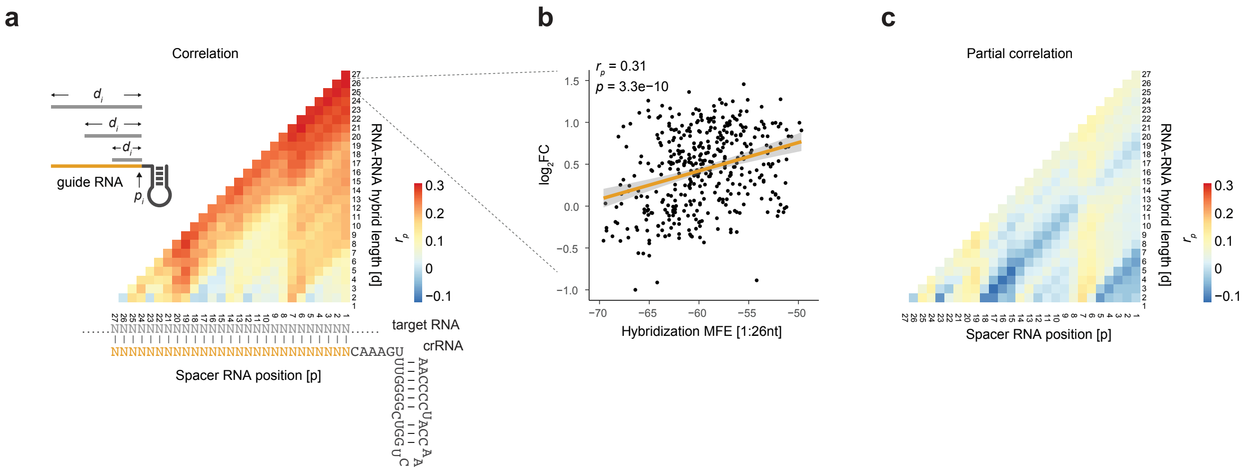


**Note Figure 3. Hybridization energy between guide RNA and target site sequence.** (**a**) Pearson correlation coefficient ($r_p$) of the observed $log_2FC$ and the hybridization minimum free energy (MFE) of guide RNA nucleotide position *p* over the distance *d* to the position *p+d* with its cognate target sequence for all perfectly matching guide RNAs. (**b**) Example scatterplot of (*a*) showing the correlation relationship between the observed $log_2FC$ and the hybridization MFE of guide RNA nucleotides 1-26 and its cognate target sites for all perfectly matching guide RNAs. (**c**) As in (*a*), but using partial correlations controlling for the crRNA MFE as shown in **Supplementary Fig. 6a**. The same $r_p$ scale is used for panels (*a*) and (*c*).

### Target site nucleotide context

Beyond guide RNA nucleotide composition, we wondered if the context features of the guide RNA target site affected target knock-down. By correlating the observed guide RNA $log_2FC$ with the nucleotide probabilities across windows around target sites, we detected a strong negative impact of high 'C'-context directly at the target site (**Note Fig. 4**). However, this may be confounded by the high guide RNA 'G'-content and its role in crRNA folding (**Supplementary Fig. 6**). Indeed, using partial correlation to account for the crRNA MFE diminished the negative correlation strength (**Note Fig. 4**). Outside the direct target site, we noticed that high 'U'-content upstream (5') to the target site positively correlated with target knock-down, which is consistent with previous reports of higher nuclease activity in 'U'-rich contexts [3,4] (**Note Fig. 4**). In order to understand if the observed upstream U-context is generalizable or targeting position specific, we generated a GFP reporter plasmid that allowed for changing the nucleotide context upstream of a perfect match target site. We designed a 52mer oligonucleotide lacking uridines and optimized to minimize predicted RNA secondary structures. We cloned this and 52mer oligonucleotides with 3 or 6 uridines at various positions into the GFP-reporter plasmid and tested the upstream uridine context effect on target knock-down. Each reporter was targeted directly downstream of the introduced oligo, or with a non-targeting guide. This was done, because the introduced uridines could potentially act as cis-regulatory elements and recruit RNA binding proteins and thus influence target RNA stability independent of the Cas13 protein [5]. We did not observe a significant position dependent effect of the upstream (5') uridines (**Note Fig 4b-c**), suggesting that the effect may be target site specific, driven by additional downstream U content or too weak to be assessed in this experiment.

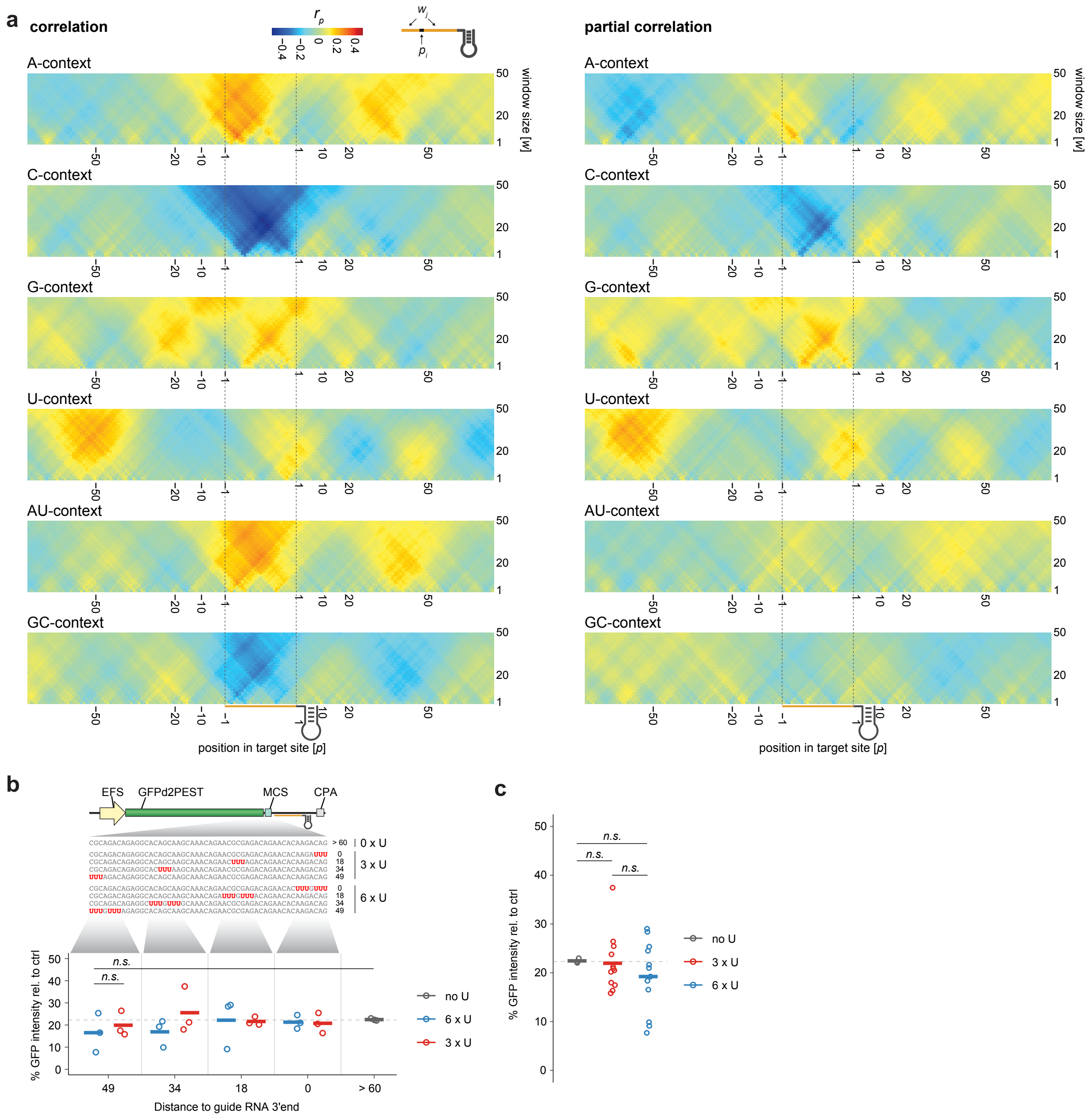

**Note Figure 4. Local target sequence context of GFP transcript targeting guides.** (**a**) Heatmaps depicting the Pearson correlation coefficient ($r_p$) between the local target nucleotide-contexts (A, C, G, U, A|U and G|C) and observed $log_2FC$ relative to guide RNA match positions. We performed a grid-search correlating the observed crRNA efficacies with the summarized target nucleotide density across a window of 1 nt up to 50 nt at every point 75 nt 5' of the target site to 75 nt 3' of the target site. (*left*) direct correlation, (*right*) partial correlation controlling for the crRNA MFE as shown in **Supplementary Fig. 6**. (**b**) Target knock-down comparison varying the upstream uridine context (position and number of uridine residues) relative to the guide RNA target site. The 60 nucleotides upstream sequence contained either 0, 3 or 6 uridines at positions 49, 34, 18 or 0 nucleotides upstream of a 26mer guide RNA. *Rfx*Cas13d-

NLS expressing cells were co-transfected with plasmids delivering the crRNA and with a GFP-encoding reporter plasmid. Shown is the percentage of mean fluorescence intensity reduction of cells transfected with a GFP-targeting guide relative to a non-targeting guide. The bar represents the mean of three replicate experiments. (**c**) As in *b* but split by number of uridines within the upstream 60 nucleotides. Differences between 0, 3 and 6 Us at each individual position, as well as summarized across all positions were not significant using one-tailed t-test.

### Target site accessibility

We also assessed whether the target site accessibility influences knock-down by correlating the observed guide RNA efficacies with the target site accessibility. Here, we define target site accessibility as the probability that the target RNA (in this case, GFP mRNA) is unpaired. We found a weak positive correlation with increased target site accessibility centered on the 3'-end of the spacer RNA (**Note Fig. 5**) reminiscent of target-RNA accessibility preferences shown for Cas13b [6].

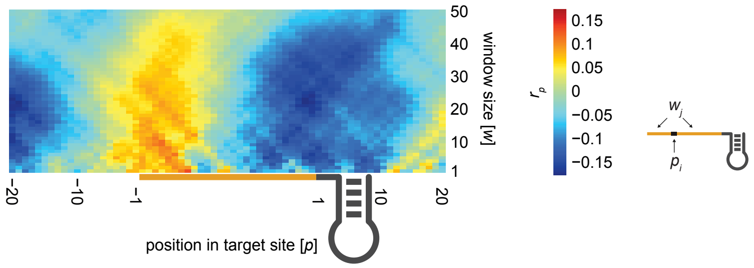


**Note Figure 5. Target site accessibility.** Heatmap depicting the Pearson correlation coefficient ($r_p$) between the local target site accessibility ( = $\log_{10}$(unpaired probability)) and the observed $\log_2$FC relative to guide RNA match positions. We performed a grid-search correlating the observed guide RNA efficacies with the unpaired probability in a window (*w*) of 1 nt up to 50 nt at every point 20 nt 5' of the target site to 20 nt 3' of the target site.

### On-target model feature collection

Based on our analyses above, we determined the position and window-size with the best correlation to the observed guide RNA enrichments for each feature (**Note Fig. 6**). A full list of all features evaluated in the on-target model based in the GFP-tilling screen data can be found in **Supplementary Table 1**.

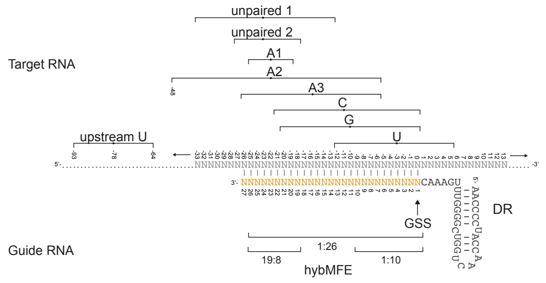


**Note Figure 6. Overview of guide RNA and target RNA feature windows**. For guide RNA features nucleotide 1 defines the guide match start site (GMSS) being the most 5' guide RNA base matching the target RNA. Nucleotide 2 relative to GMSS is the subsequent base (moving in the 5' to 3' direction) in the guide RNA and so on. For target RNA features, we denote the target nucleotide opposite to the GMSS as nucleotide 0. Moving in 5' to 3' direction target RNA nucleotide -1 is upstream to the GMSS and pairs with guide nucleotide 2, while target RNA nucleotide +1 is downstream of the target site and so on.

### Supplementary Note 2

### Features of Cas13d targeting from combined tiling screens

For the assessment of crRNA and target RNA features, we considered 2,918 perfect match guides that target coding regions across four genes: GFP, CD46, CD55, CD71 (see **Fig. 1d** and **Fig. 3a**). Compared to the 399 perfect-match guide RNAs from the GFP screen alone, this represents a 7.3-fold increase in data points. To make the screens more comparable, we scaled the $\log_2$FC of each screen independently. We preferred scaling each dataset over assigning ranks between 1 and 0 in order to maintain relative guide strength differences between screens, under the assumption that the strongest guide in screen *A* may not be as strong as the strongest guide in screen *B*. Here, each feature is represented across all 4 screens. In addition to this combined analysis, analyses for each independent screen can be found on our *gitlab* repository (https://gitlab.com/sanjanalab/cas13).

### Anti-Tag

Using all perfect match guide RNAs, we did not find evidence for the presence of an anti-tag sequence[1] for *Rfx*Cas13d suggesting that anti-tags may not be found in all Type VI CRISPRs or contribute only marginally compared to other features (**Note Fig. 1**).

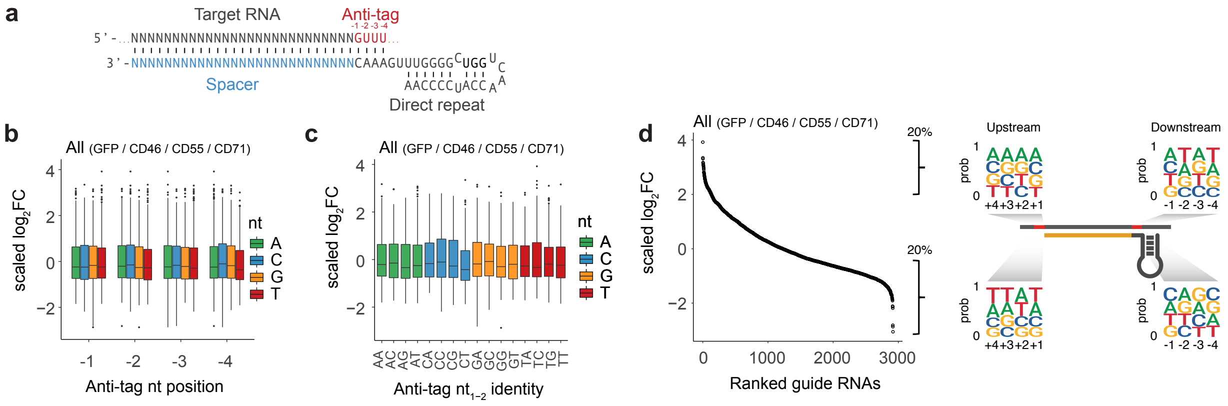


**Note Figure 7. Combined screen shows no evidence for the presence of an *Rfx*Cas13d anti-tag.** (**a**) Anti-tag position in target RNA. (**b**) $\log_2$FC of all perfect match guide RNAs ($n$ = 2,918) segregated by anti-tag nucleotide identity at anti-tag positions 1 through 4. (**c**) Same as in (*b*), but segregated by anti-tag di-nucleotide identity at anti-tag positions 1 and 2 (**d**) (*left*) $\log_2$FC-ranked guide RNAs highlighting the anti-tag sequence for the 4 least efficient guides. (*right*) Position Weight Matrices (PWMs) depicting the positional nucleotide probabilities of either the top 20% or bottom 20% $\log_2$FC-ranked guides upstream and downstream (i.e. anti-tag) relative to the guide RNA match position. (The 20% selection was applied to each screen separately to ensure equal contribution)

### Nucleotide preferences

Next, we tested whether position-based nucleotide preferences exist within the guide RNA target sequence or nearby nucleotides by comparing the nucleotide composition of the top 20% to all perfectly matching guides across all four screens, similar to previous approaches assessing Cas9 guide preferences [2]. The increased number of data points uncovered clear nucleotide preferences (**Note Fig. 2a**). The top enriched guides showed preferences for G-bases at guide nucleotides 19 –

21 (with position 1 defined as the most 5' nucleotide in the guide RNA that matches the target RNA). C-bases were favored at positions 15 – 16. Interestingly, the enrichment of G and C bases surround the center of the critical seed region at position 18 (see **Fig. 1f**). Moreover, we observed before that increased GC-content surrounding mismatches at position 18 correlated with the relative decrease in guide efficiency (delta $\log_2$FC). This suggested that increased high GC-content may ameliorate the effect size of mismatches in the seed region (compare **Supplementary Fig. 5**). There was also a mild enrichment for A- and U-bases in the first half of the guide RNA.

We correlated guide RNA nucleotide probabilities with the observed guide enrichment. In the GFP screen data alone we found that high 'G'-content in the guide RNA had a strong negative impact. This impact was reduced when taking all four screens into account (**Note Fig. 2b-c**). The guide RNA GC-content indicated a local optimum around 50% with lower guide efficiency when the guide adopts lower or higher GC proportions.

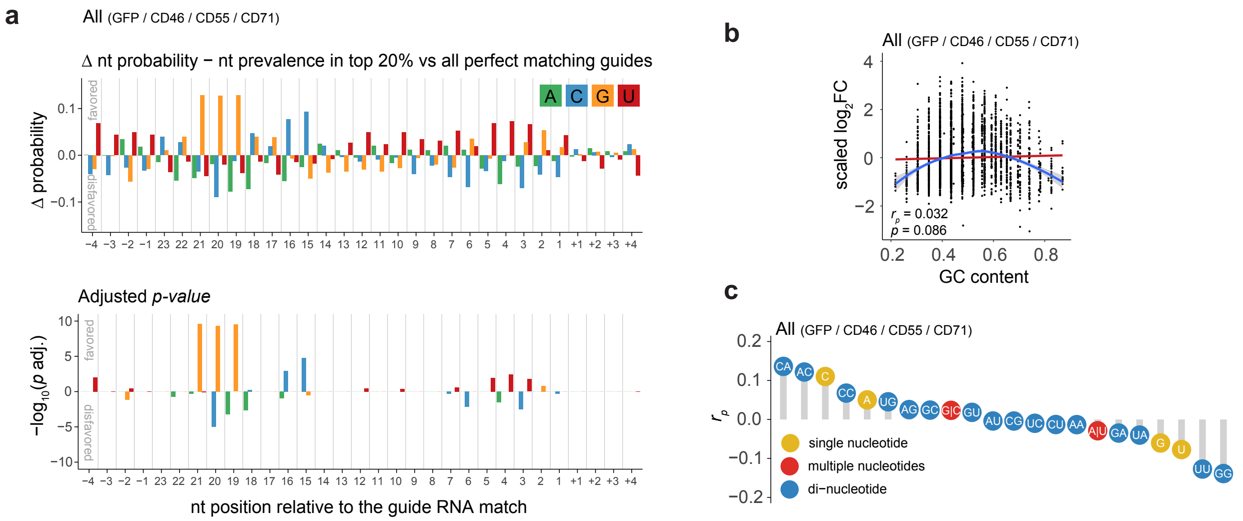


**Note Figure 8. Cas13d guide RNA nucleotide preferences influence guide RNA efficacies across all screens.** (**a**) (*top*) Effect-size (delta nucleotide probabilities) and (*bottom*) Bonferroni-corrected *p*-values of observing the conditional probability of a guide in the top 20% under the null distribution examined at every position including the 4 nucleotides upstream and downstream of the guide RNA target site. The *p*-values were calculated from the binomial distribution with a baseline probability estimated from the full-length mRNA target sequence all perfect match guide RNAs. The top 20% were selected for each screen separately to ensure equal contribution. (**f**) Scatterplot depicting the guide RNA $\log_2$FC (*y*-axis) and guide RNA GC-content as a fraction of guide length. The red line indicates the linear relationship between both values. The blue line indicates a LOESS fit and has a local optimum around a GC-content of 0.5. (**b**) As in (*e*), but showing the Pearson correlation coefficient ($r_p$) between guide RNA $\log_2$FC and all guide single nucleotides, di-nucleotides, and G|C and A|U-content.

### crRNA folding

Analyzing the GFP screen alone, we found previously that the predicted crRNA folding minimum free energy (MFE) of perfect match guides correlated positively with guide RNA efficacy (see

**Supplementary Fig. 6a**). Low MFE values were associated with low guide RNA efficiencies suggesting that stable crRNA folds may hinder crRNA utilization by Cas13d. Extending this analysis to perfect match guide RNAs of all screen, we observed an overall decrease in the correlation between crRNA MFE and guide efficiency (**Note Fig. 3**). However, low MFEs still associated with low guide RNA efficiencies. Additional predicted G-quadruplex structures were not observed in the CD46, CD55 and CD71 screens.

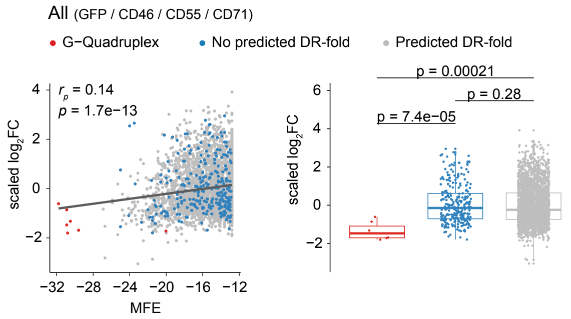


**Note Figure 9. Proper folding of the direct repeat affects crRNA targeting efficacy.** (a) Scatterplot showing the scaled guide RNA $log_2$FC versus the predicted crRNA secondary structure minimum free energy (MFE). The Pearson correlation coefficient ($r_p$) is nearly unchanged ($r_p$ = 0.12) when MFEs of G-quadruplex-forming crRNAs are ignored.

### Guide RNA – target RNA hybridization

We next tested whether guide-target hybridization can contribute to guide RNA efficacy when integrating data from all four screens. Unlike for the GFP screen alone, we found that the overall hybridization energy between the full-length guide RNA and target sequence correlated less (**Note Fig. 3a**). Instead, the hybridization energies of sub-fragments contributed differentially to the overall guide-target interaction. The hybridization energy between the 12 nucleotides from guide position 3 to 15 and the cognate target site showed a slight positive correlation (**Note Fig. 3a-b**). Hybridization energies covering the 9 nucleotides from guide position 15 to 23 correlated negatively with the knock-down efficiencies (**Note Fig. 3a-b**). Unlike for the GFP screen analysis before, these sub-structures were still present when controlled for the crRNA folding energies using partial correlations (**Note Fig. 3c**).

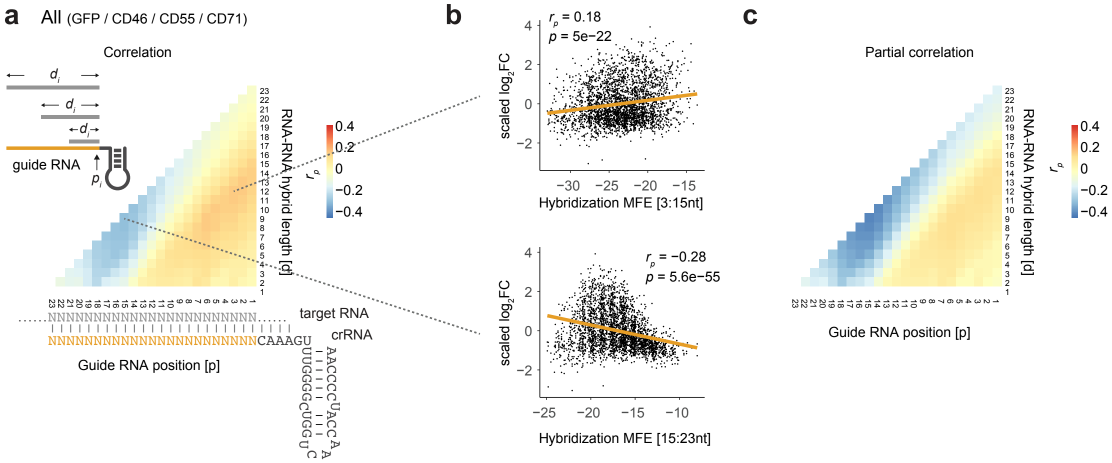


**Note Figure 10. Hybridization energy between guide RNA and target site sequence.** (**a**) Pearson correlation coefficient ($r_p$) of the observed scaled $\log_2$FC and the hybridization minimum free energy (MFE) of guide RNA nucleotide position *p* over the distance *d* to the position *p+d* with its cognate target sequence for all perfectly matching guide RNAs. (**b**) Example scatterplot of (*a*) showing the correlation relationship between the observed scaled $\log_2$FC and the hybridization MFE of guide RNA nucleotides 3-15 (top) and 15-23 (bottom) and their cognate target sites for all perfectly matching guide RNAs. (**c**) As in (*a*), but using partial correlations controlling for the crRNA MFE as shown in **Note Fig. 3**. The same $r_p$ scale is used for panels (*a*) and (*c*).

### Target site accessibility

We also assessed the target site accessibility for all screens and correlated observed guide RNA efficacies with the target site accessibility. Here, we define target site accessibility as the probability that the target RNA is unpaired. We did not find a strong relationship between the probability of the target sequence being unpaired and the observed knock-down strengths (**Note Fig. 5**). Similar to the GFP screen alone, we find a weak positive correlation with increased target site accessibility centered on the 3'-end of the spacer RNA. We also recapitulate the observed nucleotide preferences with higher GC-content centered around seed nucleotide 18, which is surrounded by higher AU-content (compare **Supplementary Figure 10a-b**). Higher AU-content translates to increased accessibility, while higher GC content suggest local secondary structures.

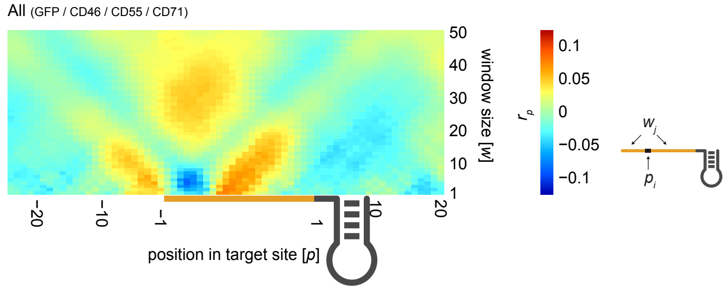


**Note Figure 11. Target site accessibility.** Heatmap depicting the Pearson correlation coefficient ($r_p$) between the local target site accessibility ( = $\log_{10}$(unpaired probability)) and the observed $\log_2$FC relative to guide RNA match positions. We performed a grid-search correlating the observed guide efficacies with the $\log_{10}$-transformed unpaired probability in a window (*w*) of 1 nt up to 50 nt at every point 20 nt 5' of the target site to 20 nt 3' of the target site.

### On-target model feature collection

Based on our analyses across all four tiling screens, we determined the position and window-size with the best correlation to the observed guide RNA enrichments for each feature (**Note Fig. 6**). For the RNA target site accessibility we chose the entire target site as a window instead of the weak positive correlation that correlated with the U-context in in that region (from nucleotide 1 – 23 with position 1 defined as the most 5' nucleotide in the guide RNA that matches the target RNA). A full list of all features evaluated in the on-target model based in the GFP-tilling screen data can be found in **Supplementary Table 2**.

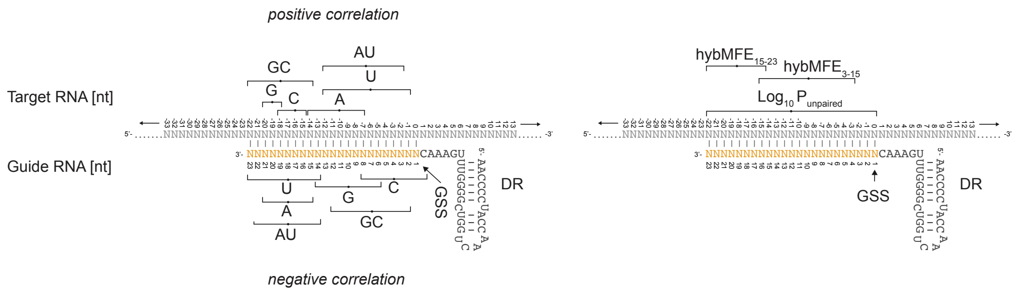


**Note Figure 12. Overview of guide RNA and target RNA feature windows**. For guide RNA features nucleotide 1 defines the guide match start site (GMSS) being the most 5' guide RNA base matching the target RNA. Nucleotide 2 relative to GMSS is the subsequent base (moving in the 5' to 3' direction) in the guide RNA and so on. For target RNA features, we denote the target nucleotide opposite to the GMSS as nucleotide 0. Moving in 5' to 3' direction target RNA nucleotide -1 is upstream to the GMSS and pairs with guide nucleotide 2, while target RNA nucleotide +1 is downstream of the target site and so on. Selected features with either positive or negative correlation are denoted with the subscript 'max' or 'min', respectively, in **Supplementary Table 2**.
